# Dynamics of the end-of-life phase explained by the saturating removal model

**DOI:** 10.64898/2026.04.20.719588

**Authors:** Naveh Raz, Glen Pridham, Michael Rera, Kenneth Rockwood, Uri Alon

## Abstract

End of life is characterized by a phase of rapid physiological decline and high morbidity, phenotypically observed as the “Smurf” phase in *Drosophila*, metabolic end-of-life dysregulation in mice, and end-stage frailty in humans. Existing two-phase aging models often conceptualize this end-of-life phase as a discrete biological state. Here, we demonstrate that a continuous stochastic model of damage accumulation, the saturating removal (SR) model, captures these multi-species morbidity dynamics. By defining the end-of-life phase as a stochastic crossing of a sub-lethal damage threshold, the SR model accurately reproduces empirical end-of-life dynamics across flies, mice, and humans. The model predicts a surprising temporary reduction in hazard shortly after entering the end-of-life phase consistent with empirical data in all three organisms. It also correctly predicts a shortening twilight phenomenon where the mean duration of the end-of-life phase decreases the later its onset. We conclude that end-of-life dynamics are consistent with universal features of a driver of aging crossing a threshold for end-of-life morbidity and then a threshold for death.

## Introduction

The aging field is data rich, and can benefit from unifying theoretical models^1–5^. Such mathematical models can help design new experiments and interventions, and translate findings between model organisms and humans.

Some models, like the gamma-Gompertz equations, are commonly used to fit mortality data but have no mechanistic basis^6^. Other classical models such as Streller Mildvan^7,8^, reliability^9^ and network models^10^ are suggestive but difficult to connect with specific molecular and cellular components. There is a need for mechanistic models that connect multiple quantitative patterns of aging with molecular and cellular processes^1,11^.

One of the most extensively validated mechanistic models of aging is the saturated removal model presented by Karin et al^12^. The model was developed based on longitudinal measurements of senescent cells in mice, and has been found to also explain longitudinal membrane damage dynamics in bacteria^13^. It postulates a damage *x* that drives aging - although it is agnostic to the nature of *x*, and can thus apply to different organisms with different molecular drivers^1,12,14^. Several plausible candidates for *x* in different organisms have been suggested^1,12^.

The SR model tracks damage *x* using a stochastic differential equation, in which production rate rises linearly with age, and damage is removed by a process that saturates at high damage. Death occurs when *x* crosses a death threshold *X*_*c*_.

The SR model captures the Gompertz law in humans and other organisms, and the Weibull hazard in invertebrates^1,12,14^. It suggests that evolution has tuned lifespan changes between species mainly by varying the damage production rate^14^. The model was used to revise estimates of the heritability of human lifespan^15^, and to explain senescence levels in parabiosis experiments in mice^16^.

An important development was to add to the SR model the ability to describe age-related disease states. Katzir et al postulated that disease onset occurs when *x* crosses a disease-specific threshold *X*_*d*_ in susceptible individuals. This simple extension quantitatively explained the exponential rise in incidence of hundreds of age-related diseases from medical records^17^. The same definition of a disease threshold was used by Yang et al to discover which longevity interventions compress morbidity in invertebrates and mice^18^.

The power of mathematical models is that they can be refuted by data, allowing improvement of understanding. It is therefore of interest to find additional aging phenomena to test the saturated removal model and its disease-threshold extension.

Here we test the SR model against the dynamics of end-of-life phases in different organisms. Rera and co-authors^19–21^ showed that fruit flies fed with a blue dye enter a whole-body blue phenotype a few days before death called the Smurf phase. Rera et al also found evidence for an analogous terminal phase in mice, Zebrafish and *C. elegans*^21,22^. This has been modeled as a two-step process^19^ : In the two-step model, individuals have a linearly rising probability with age to become Smurf - the damage accumulation phase, and then a constant probability per unit time to die - the dying phase. Clinical observations in humans show a qualitatively similar end-of-life phase known as end-stage frailty, where physiological reserves are depleted and risk of adverse outcome increases sharply^23^.

We asked whether the SR model can capture the dynamics at the end of life. We find that modelling the end-of-life phase as a crossing of a sublethal threshold captures the dynamics of the Smurf phase in flies and mice, and the dynamics of end-stage frailty in humans. We show that in the three organisms the end-of-life phase has common features, including an initial drop in hazard after the end-of-life phase onset, and a reduction in the mean end-of-life-phase duration the later it is entered. We conclude that end-of-life dynamics are consistent with universal features of SR model damage crossing a sublethal threshold near the end of life.

## Results

### SR model with threshold for Smurf onset captures the two-phase dynamics of fruit fly aging

The SR model is a stochastic differential equation in which production rises linearly with age, η*t*, removal saturates with damage,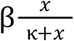, together with additive noise (**Fig 1a**). Death occurs when *x* crosses the death threshold *X*_*c*_. Simulations of the model show stochastic trajectories of damage that fluctuate around an accelerating trend.

**Figure 1.**
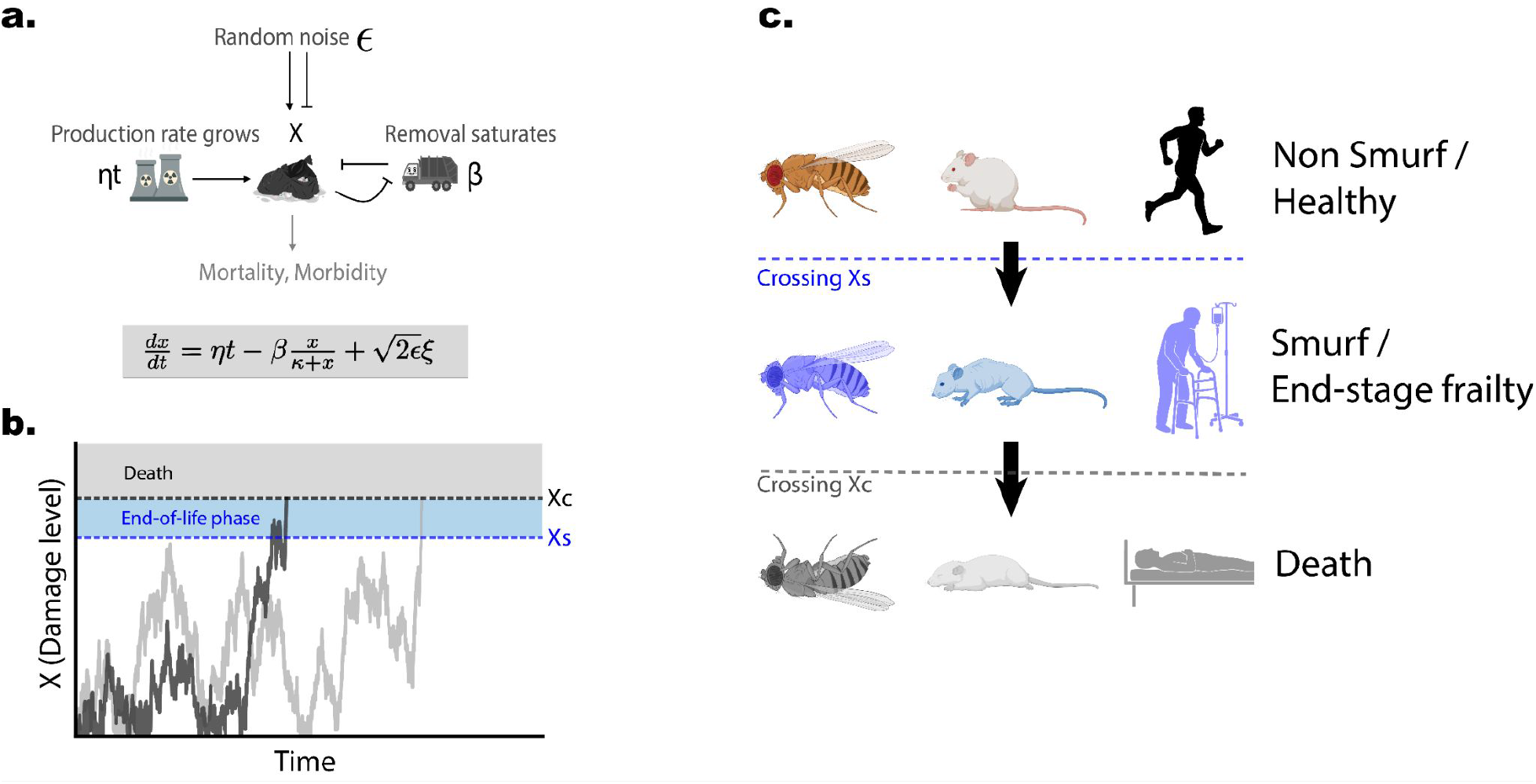
SR model and Smurfness/end of life schematics. **a**. SR model describes the dynamics of an uncharacterized damage *x* causal for aging as a balance of production, removal and noise. The damage gives rise to both mortality and morbidity. **b**. Simulations of many individuals show noisy damage trajectories. End-of-life phase onset occurs at the first crossing of the end-of-life threshold *x* > *X*_*s*_, death occurs when damage crosses the death threshold *x* > *X*_*c*_. **c**. Different organisms transition to end-of-life phase and then die. *Drosophila* flies are fed a nontoxic blue food dye. Before death their guts become permeable to the dye and they turn blue (Smurfs). In mice end-of-life (Smurf) phase is defined as a change in dynamics of several physiological functions and in humans the end-of-life phase is defined as end-stage frailty.

To model the end-of-life phase, we postulate that it begins when damage *x* crosses an end-of-life threshold *X_s_* **(Fig 1b**). Thus every simulated individual has a “non-end-of-life” phase followed by an end-of-life phase, and ultimately death (**Fig 1c**). While the underlying systemic damage fluctuates stochastically and may temporarily drop below *X*_*s*_, we define the onset of the end-of-life phase as the *first* crossing of this threshold.

Once an individual enters this phase, they remain in it until death. This continuous classification aligns with the irreversible nature of the Smurf transition observed in flies and mice. All individuals go through the end-of-life phase, even if only briefly.

In humans the end-of-life phase is less clearly defined, and we use a frailty index^23,24^ as a proxy as described below, despite low-probability cases of significant improvement in frailty index after high frailty (as detailed in **Supplementary Note 1**). From a mathematical perspective, the SR model does not strictly enforce this irreversibility and our analysis focuses strictly on the dynamics following the initial entry into this phase. Therefore, our findings hold regardless of whether the underlying biological state occasionally reverses in humans.

We fit the SR model parameters to the survival curve of *Drosophila drs*GFP as published by Rera et al^19^ (**Fig 2 a,b**). We find good fits to the hazard (*R*^2^ = 0. 96) and survival curves (*R*^2^ = 0. 99). We then identify a Smurf threshold *X*_*s*_ that best fits the non-Smurf survival curve (**Fig 2c**). As expected, this threshold *X*_*s*_ = 3. 4 is lower than the death threshold *X*_*c*_ = 4. 1. The model also captures the distribution of lifetimes in the Smurf phase that drops roughly exponentially (**Fig 2d**) as previously described in Tricoire and Rera.

**Figure 2.**
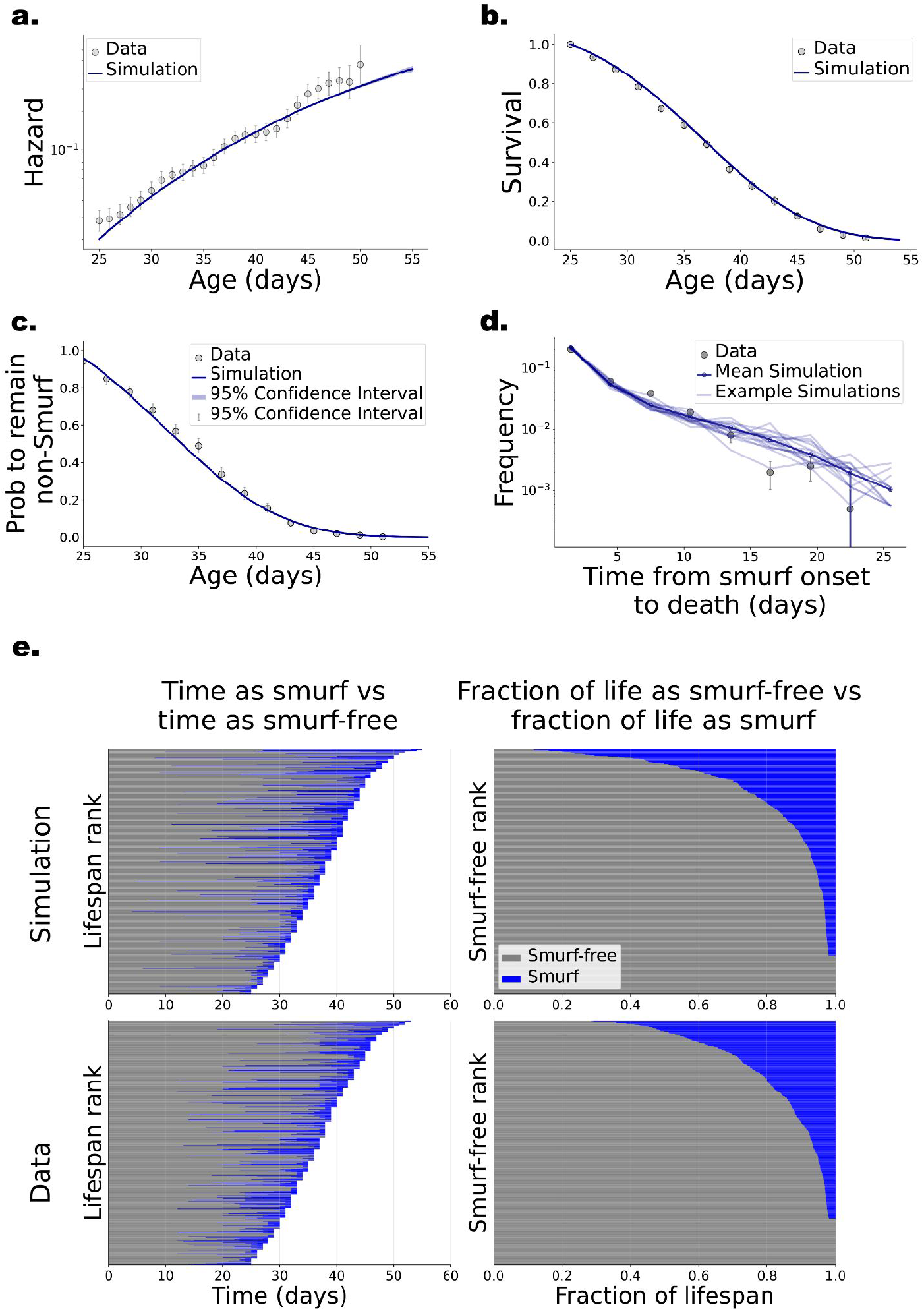
SR model captures Smurf dynamics in *Drosophila*. **a+b**. Death hazard rates and survival curves, measured vs simulated. **c**. Smurf-free survival curves as measured and simulated show the probability to remain non-smurf as a function of age. **d**. Distribution of time until death from the time of smurf onset. **e**. Smurf and smurf-free portion of life-span for each individual. Same simulation (top) and data (bottom) as a function of age (left, ranked by lifespan) or proportion of lifespan (right, ranked by smurf proportion), individuals that die before age 25 days are excluded (see methods and SI). In d. we binned the data every 3 days, error bars are std calculated via bootstrapping. Error bars are 95% CIs. N data = 668, simulation parameters are 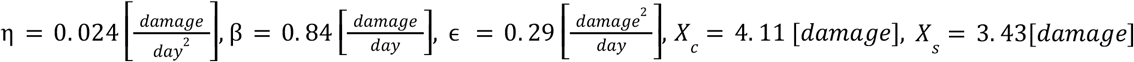

### SR model correctly predicts that mean survival time in Smurf phase decreases with onset age

One behavior in which the SR model disagrees with the two-step aging model of Tricoire and Rera^19^, is in the average survival time in the Smurf phase. The two-step model postulated that the mean survival time is independent of the age of onset of the Smurf phase^19^. In contrast, the SR model predicts that the mean Smurf survival time decreases with Smurf onset age. The reason for this drop is that older onset means a higher damage production rate η*t* and hence faster demise on average.

We therefore re-analyzed the mean survival time from the data from Rera et al ^19^. We confirmed that the average Smurf survival time drops from about 5 ± 1 days in flies with early Smurf onset (25 days) to about 1 ± 0. 5 days in flies with late Smurf onset (50 days). Cox regression indicates that for every day that Smurf onset is delayed, the hazard of death following onset increases by roughly 4%(95% CI: 2%-6%, p < 0.005). Results for the simulated data were similar, showing an increase in the hazard of roughly 5% per day, and a different model by Breuil et al found 3.2%^25^, both are within error bars of the data. We conclude that the SR model prediction agrees with the measured mean Smurf survival time versus Smurf onset age, without fitting any new parameter (**Fig 3**).

**Figure 3.**
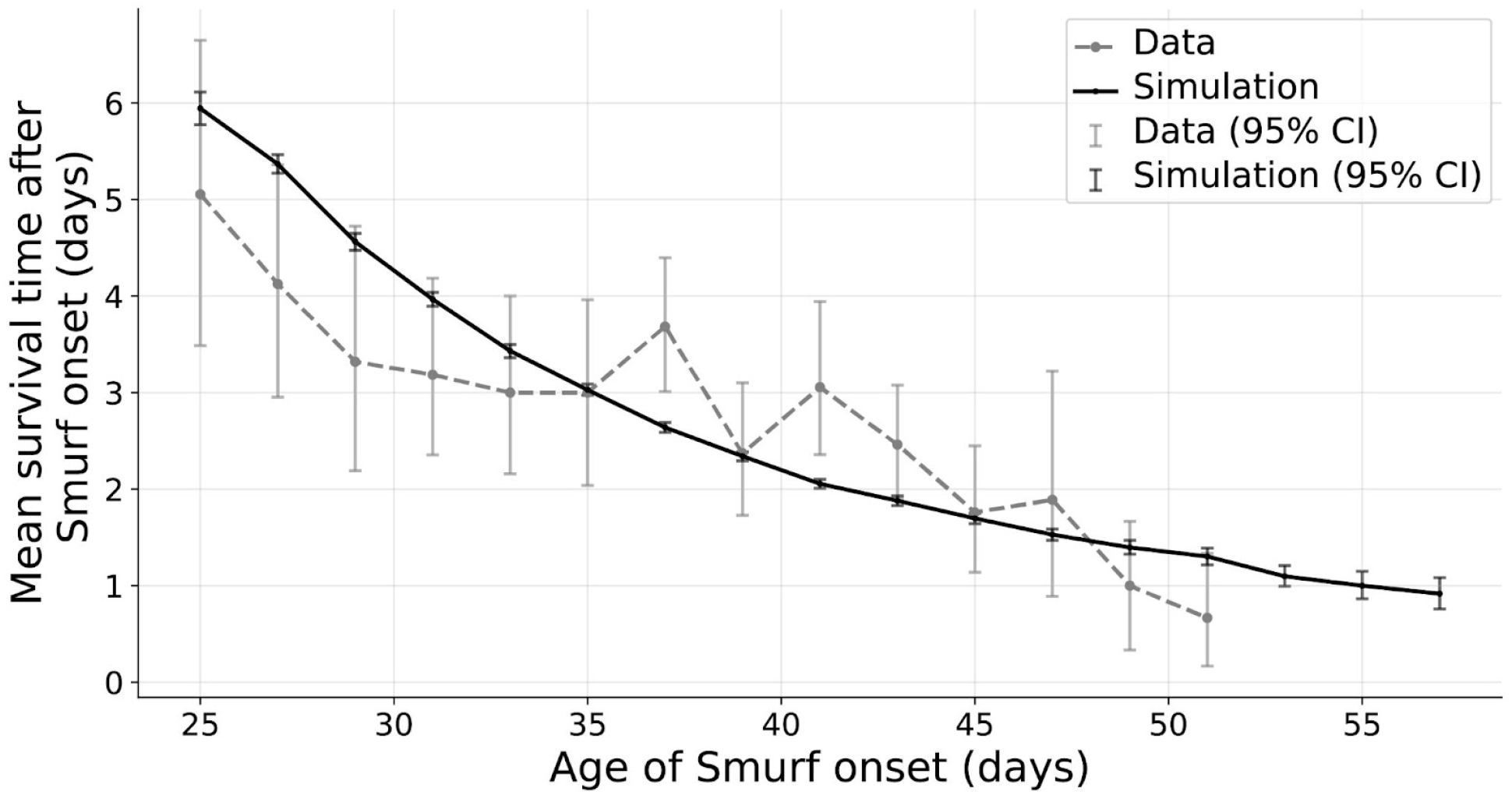
The SR model captures the reduction of Smurf mean duration with onset age in fruit fly. Simulation and data show that mean time spent as a Smurf is shorter the later the Smurf onset, in contrast to what was previously described using the median Smurf durations.

In contrast to the average Smurf lifespan, the median Smurf lifespan shows weaker dependence on age of onset. This is because many flies die in the first Smurf days, but at a young age many Smurf flies survive for longer times, a tail that is picked up by the average but less so by the median (**supplementary note 2**). Of note, Rera et al originally reported the median Smurf lifespan as 3.1 days^20^ and Tricoire and Rera as 2.04 days^19^.

To further validate the model, we fitted the SR parameters to survival data from various *Drosophila* genotypes. In agreement with the empirical observations reported by Tricoire and Rera^19^, the SR model accurately predicts comparable median Smurf durations across these different strains (**supplementary Note 3**)

We conclude that the Smurf phase is consistent with a threshold-crossing phenomenon in the SR model of aging.

### Mouse end-of-life phase captured by the SR model

We next turn to mice, and consider the experiments of Cansell et al, which defined an analog of the Smurf phase in mice of two different strains, AKR/J and C57BL6/J. The Smurf phase was defined using the dynamics of five longitudinal physiological markers, specifically glycemia, body weight, food intake, fat mass and body temperature. All of these markers showed a change in dynamics near the end of life. The Smurf phase was defined for each mouse using a change point model, specifically as the time by which three of the markers crossed their change point. The mean Smurf duration was two weeks in AKR/J and three weeks in C57BL6/L mice.

We applied our modeling to this mouse dataset, defining an end-of-life threshold for each strain and sex. We find that the SR model agrees very well with the survival curves and the time-to-Smurf-onset distribution (**Fig 4a-f**). The model also agrees well with the distribution of Smurf phase durations (**Fig 4 g-i**).

**Figure 4.**
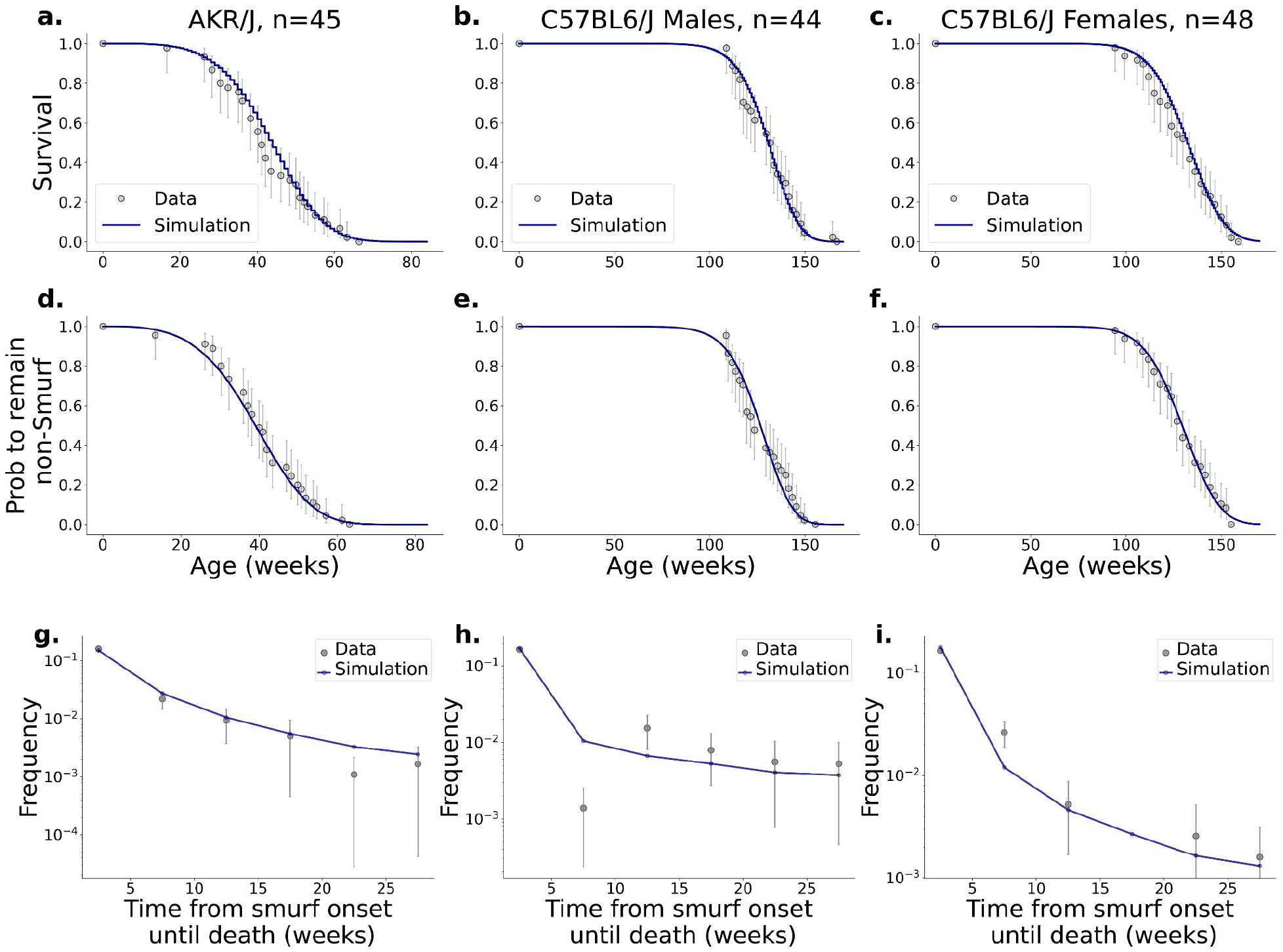
The SR model captures end-of-life phase dynamics in different mice strains. The model captures the Smurf dynamics in mice as described by Cansell et al.^22^ **a-c**. Survival curves of different mice strains. **d-f**. Smurf free survival curves. **g-f**. Distribution of time until death from the time of Smurf onset. For full fit parameters see supplementary note 4. Error bars are 95% CIs.

### The SR model captures the dynamics of the human end-of-life phase defined by the frailty index

We also considered human frailty index data to explore a frailty end-of-life phase. The frailty index is a widely-used measure, defined as the fraction of deficits that a person has from a list of deficits including clinical, pathological and functional measures^23,24^. It ranges between zero and one, and life is often incompatible with a frailty index above 0.7^23,26,27^.

We defined the end of life phase using frailty index above 0.55, defined as end stage frailty by Kim and Rockwood ^23^. A sensitivity analysis confirms that while changing this threshold alters the duration of the end-of-life phase, the core dynamics remain consistent, though model fits are better at the 0.55 threshold then for lower thresholds (**supplementary note 13**). The end of life phase averages 1.3 years in men and 1.7 years in women.

We find that the SR model captures end-stage frailty dynamics with a threshold model, where the end-of-life threshold is 97% of the death threshold for males, and 96% for females (**Fig 5**). The SR model captures the approximately exponential distribution of end-of-life phase duration (**Fig 5 e,f**).

**Figure 5.**
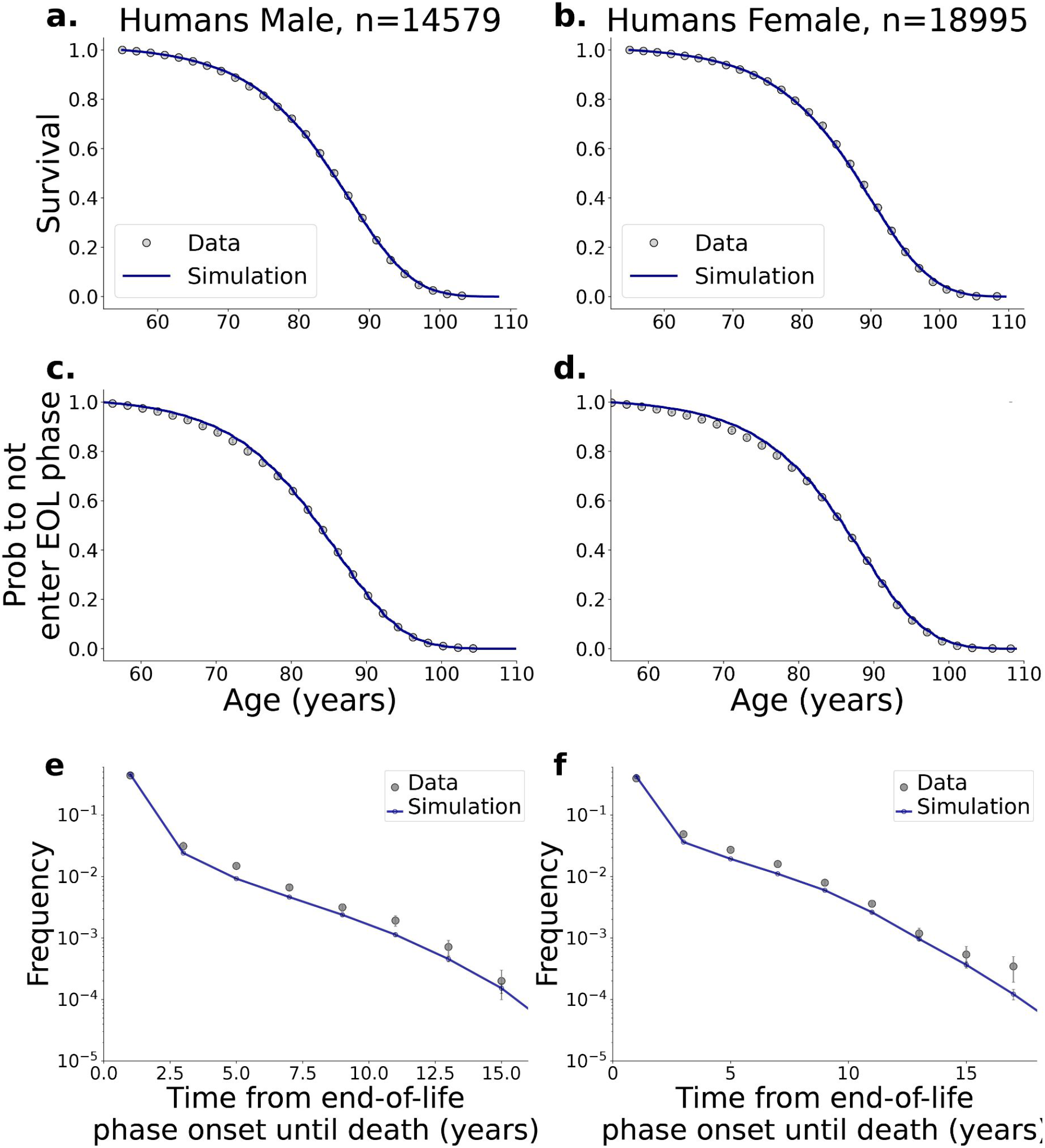
End of life dynamics in humans captured by the SR model. We defined the end of life phase as the part of life after reaching severe frailty^23^ (frailty index>0.55) **a-c**. Survival curves. **d-f**. Probability to not have not entered the end-of-life phase as a function of age. **g-f**. Distribution of time until death from the time of end-of-life onset. For full fit parameters see supplementary note 4. Error bars are 95% CIs and are smaller than the markers for a-d.

Like in flies, the model also captures a “diminishing twilight” in humans where the duration of the end-of-life phase decreases with the age of onset. See s**upplementary note 5**.

### Hazard shows a transient decline upon entry into the end-of-life phase

The SR model makes a subtle prediction for the hazard of death as a function of time in the end-of-life phase. Initially, the hazard is high because individuals have recently crossed the sublethal threshold *X*_*s*_, and thus their systemic damage level is close to the death threshold *X*_*c*_. Subsequently, the hazard declines as stochastic noise and removal processes allows systemic damage to regress toward lower levels. Notably, these individuals remain classified within the end-of-life phase even if their damage levels temporarily drop back below *X*_*s*_.

At more advanced ages, the hazard rises again because the age-dependent increase in damage production (η*t*) eventually dominates, driving damage trajectories toward the death threshold. This combination of stochastic crossing and age-driven production results in a characteristic U-shaped hazard curve in simulations (**Fig. 6 a,b**)

**Figure 6.**
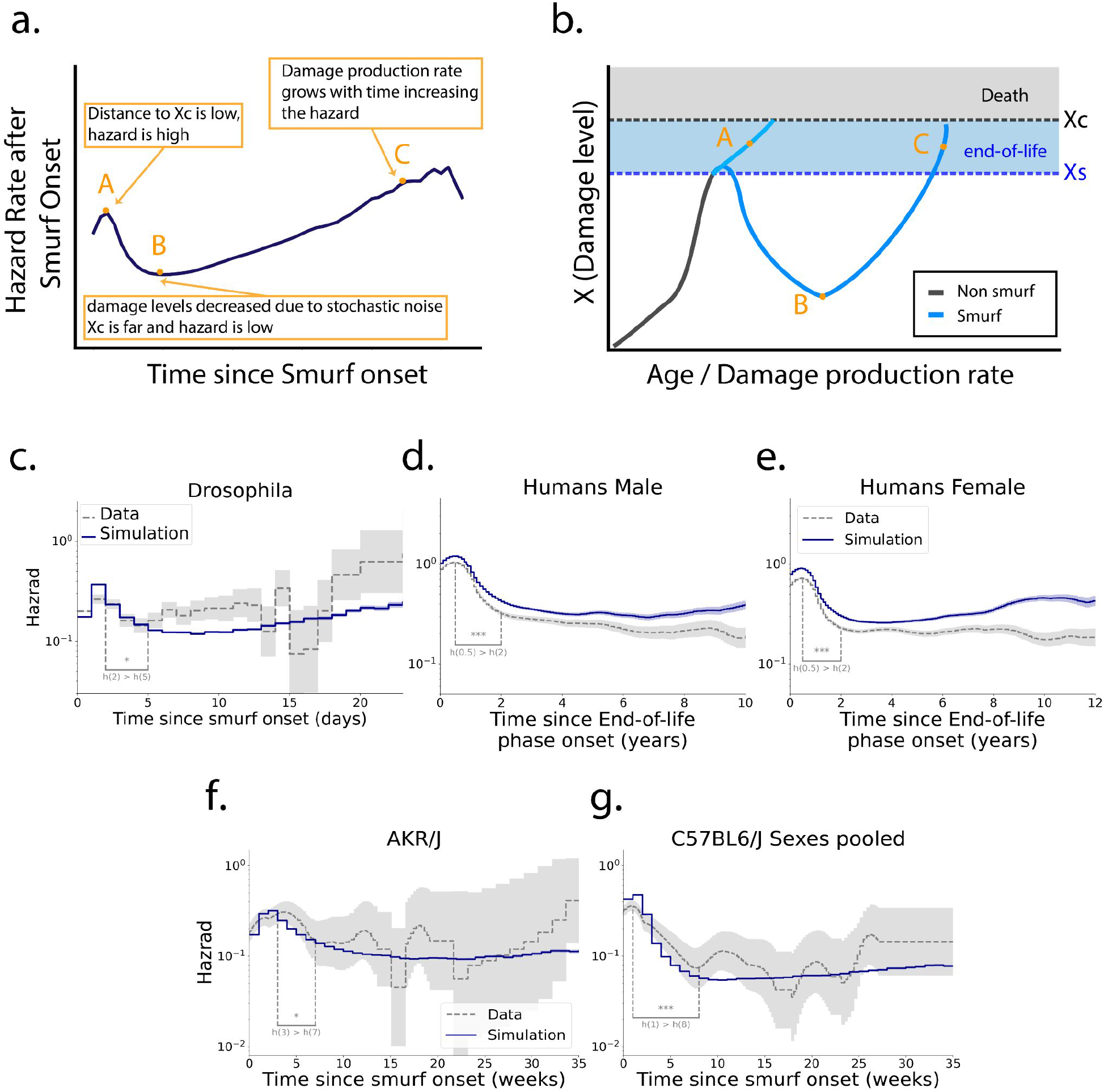
the end of life phase shows an initial decline of hazard. **a**. The SR model predicts that hazard as a function of time in the end of life phase initially drops and then rises (resulting in a U shape), because **b**. After crossing the end-of-life threshold damage is high and mortality is likely (A), then if the death threshold wasn’t crossed damage may temporarily decrease reducing the hazard (B), finally the production rate η*t* rises with age increasing the hazard again (C). The end of life phase begins after crossing the Smurf threshold and persists even if damage levels drop below it. The Simulations in *Drosophila* **c**, humans **d**,**e**, and mice **f-g**, (blue line) are consistent with an initial decline of empirical hazard in the end of life phase (dashed lines). Confidence intervals are 68%. Drosophila *h*(2 *days*) > *h*(5 *days*), *p* = 0. 02, human *h*(0. 5 *year*) > *h*(2 *years*), *p* < 0.001 (both sexes). AKR/J *h*(3 *da*y*s*) > *h*(7 *days*), *p* = 0. 02, C57BL6/J *h*(1 *da*y) > *h*(8 *days*), *p* < 0. 001. *p* values calculated via bootstrapping.

The end-of-life mortality data in flies and mice is consistent with this prediction. Empirical hazard in the end-of-life phase follows roughly a U shape pattern (**Fig 6 c-h**), as noted by Breuil et al in flies^25^. In human data, the initial decline of hazard is clearly seen (*hazard*(2 *years*) < *hazard*(0. 5 *years*), *p* < 0. 001)). The late rise is not seen in the data, potentially due to limited followup times. The rise is seen in various subpopulations stratified by age (**supplementary note 12**).

To confirm that the initial decline and this U-shape is a biological feature rather than an analytical artifact, we demonstrated that randomizing the end-of-life onset times abolishes these effects (**supplementary note 14**).

## Discussion

We find that the SR model quantitatively explains the dynamics of the end-of-life phase in flies, mice and humans, by describing it as a threshold crossing of the damage variable that also drives mortality. The SR model predicts that the average end-of-life phase duration is longer in individuals that reach it at a young age compared to those that reach it at an older age. This prediction is described using an independent modeling approach by Breuil and co-authors, and quantitatively matches data in flies, mice and humans, where the latter is described using the frailty index with end stage frailty serving as the human parallel to the Smurf phenotype during the end-of-life phase ^23^. The model also correctly predicts an initial decline of hazard after entering the end-of-life phase. The SR model has been used elsewhere as a tool to analyze cellular and molecular aging, human disease and longevity interventions - and hence its ability to explain the end-of-life phase offers a way to connect experimental models such as the fly Smurf phase to understand human morbidity and end of life stages.

It is natural to think that the risk of death should rise as a function of time spent in the end-of-life phase. It thus may seem surprising that the hazard initially declines. The temporary drop of hazard soon after the onset of the end of life phase is due to trajectories of damage that stochastically return to low levels. This is a general feature of stochastic crossing of two thresholds, and is seen in flies, mice and human data. Interestingly, an initial decline and later rise (U-shaped) hazard pattern is also seen in studies on human colorectal cancer mortality^28^ and mortality post heart transplantation^29^. An alternative explanation to the initial hazard decline is heterogeneity in the death threshold - individuals with low death threshold are more likely to die early, leaving a population with higher death thresholds which has lower hazard ^15,30^. The SR model with an end-of-life threshold relates to the model used by Katzir et al^17^ to describe the onset of age-related diseases. Both models assume that the damage that drives death also drives observable aging phenotypes – end-of-life phase here, and disease onset in Katzir et al. One difference is that Katzir et al included another parameter - the fraction of the populations susceptible to each disease. If only a fraction of the population is susceptible, as in most diseases, the incidence curve rises exponentially but then drops at very old ages. The decline does not happen if susceptibility is 100%, as it is in the Smurf phase which occurs in all flies.

The threshold crossing in the Katzir model is based on a framework in which each disease is a tipping point of a certain physiological system. The rising damage *x* perturbs the parameters of a physiological system until, at a critical damage level *X*_*d*_, the system bifurcates to a pathological behavior. For example *x* affects cancer cell proliferation and removal rates through inflammation, saturation of immune removal and other factors. At a critical level of *x*, the rate of cancer cell proliferation exceeds removal and cancer grows exponentially. Similarly, it is possible that onset of the Smurf phase is a tipping point, driven by the primary driver of aging *x*, that once tipped causes systemic changes detected thanks to the intestinal permeability phenotype^31,32^.

The observed decline of average duration of the end-of-life phase as a function of onset age is an instance of a phenomenon known as “shortening twilight“. The twilight - the span between onset of an age-related phenotype and death - shortens with the age of onset. Shortening twilight has been documented by Stroustrup et al in *C. elegans*^33^, and in *E. coli* longitudinal damage measurements^13^. Shortening twilight is a general prediction of the SR model, because onset (at any age) occurs at the same damage level *X*_*d*_, but production rate is higher at old age, and thus death is reached faster on average at old onset than in young onset from the same damage starting point. Shortening twilight is thus a test of the SR model with a pathological threshold.

The SR model makes additional predictions for future experiments regarding the end-of-life phase duration under lifespan-extending interventions. Building on the work of Yang et al^18^, the model can predict which longevity intervention would expand or shrink the end-of-life phase relative to lifespan. Interventions that extend median and maximal lifespan by the same factor, preserving the shape of the survival curve, are predicted to also expand the end-of-life phase duration by the same factor. An example is slowing the rate of damage production. In contrast, most interventions that extend median lifespan more than maximal lifespan (steepening the survival curve) are predicted to shrink the proportion of lifespan in the end-of-life phase. An example is speeding the rate of removal of *x* (which keeps the end-of-life phase similar in absolute terms and smaller relative to lifespan). This is analogous to compression of morbidity found in steepening interventions in mice and invertebrates ^18^. A related question can apply to different strains. Tricoire and Rera showed that median Smurf phase is similar in fly strains with a 3-fold difference in lifespan. Simulations of the model for different fly strains capture this as well (**supplementary note 3)**.

The main focus of this study was to characterize the end of life phase and test the SR model. Other models of aging can in principle also capture the main features of shortening twilight, initial decline in hazard in the end-of-life phase, and the death and end-of-life onset times distributions. This includes stochastic differential equation models such as Fedichev and Gruber in their stable regime^34^, but not the in their unstable regime in which dynamics are time invariant (time invariance precludes shortening twilight if the end-of-life threshold is defined by the same level of *x* = *X*_*s*_ at different ages). Other damage models such as queue theory models with constant production and time decreasing removal can also capture these features (but they do not capture the reducing CV of *x* found in data, as mentioned in Karin et al)^17,35^. Since by now the SR model has been tested with a wide range of empirical phenomena, a comprehensive comparison of different aging models across phenomena is warranted.

We find that end-of-life phases show dynamics that have universal features explained by the SR model - in which end-of-life is described as damage crossing a sublethal threshold followed by a death threshold. These universal features include a temporary reduction in hazard soon after the end-of-life phase is entered. A second feature is shortening twilight where end of life is shorter on average the later its onset. It would be fascinating to explore this in additional organisms and settings. Since the SR model can describe the effects of interventions^18^, it may help to understand the impact of future gerotherapeutics on end-of-life morbidity and survival.

## Methods

For model simulation and parameter calibration we used the same pipeline as in^14^ which we generally outline with the relevant modifications for the end-of-life state below.

### Stochastic model simulation

Simulation of mortality data was done using python. We simulated the SR model stochastic differential equation as an Itô process 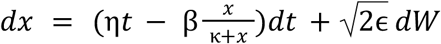 and calculated the death time of each individual as a first passage time of *X*_*c*_ and the Smurf onset as the crossing of an earlier threshold *X*_*s*_. Where relevant, survival and hazard functions were calculated using the python Lifelines library.^36^

### Bayesian estimation of parameters

We used a Markov Chain Monte Carlo (MCMC) using the emcee python library^37^ to estimate the model parameters as described in^14^. The differences included the inclusion of Smurf times in the simulation and a modified likelihood function that accounts for the Smurf times as described in **supplementary note 6**. Briefly, we selected parameters with the maximum *a posteriori* using non-informative priors for each parameter as before but extend the likelihood function to simultaneously include both smurf time and survival time. Importantly, this additional degree of freedom improved the estimation of *X*_*c*_ which previously had large uncertainty.

### Data preparation

Drosophila *drs*GFP data is from Tricoire & Rera^19^. N before filtering was 1159 mated females, monitored individually approximately once every 24h. After filtering 668 individuals remained. Flies were transferred to the blue medium on day 12^19^, we therefore allowed for an adjustment period and filtered the data for the drsGFP flies at day 25. See **Supplementary note 7**.

Mice data is from Cansell et al^22^. The Mice were sampled every 1-2 weeks. Sample sizes were N = 45 (AKR/J,Females), 44 (C57BL6/J Males), 48 (C57BL6/J Females).

Human data are from the Health and Retirement Study (HRS)^38^. We used a threshold of FI>0.55 inline with the definition of end-stage frailty by Kim and Rockwood^23^. The effects of varying this threshold on survival curves and model likelihoods are detailed in **supplementary note 13**.

The frailty index is the average of 41 binary self-reported multi-domain health variables, including physical limitations, signs, symptoms and diagnoses, as summarized in **supplementary note 8**^39^. We used waves 6-15 (2002-2020) of the RAND preprocessed files; waves are measured every 2 years. This included 34672 individuals with 189096 visits; median age: 67 (inter-quartile range: 59-76). The frailty index ignored missing data; 1% of data were missing, the least commonly measured variable was difficulty climbing several flights of stairs (6% missing).

For Mice and human data, we have regularly sampled data rather than transition times and so we assumed the transition to end-of-life phase occurred with equal probability during the interval between the first observation of end-of-life phase (ie, Smurfness or FI>0.55) to the previous observation. In cases where the individual died without observing the end-of-life state, we assumed the transition occurred between the time of death and the last observation before death.

### Cox regression

Cox regression for dependence of age of Smurf onset was performed using python lifelines library^36^.

### Fractional binning to account for EOL onset uncertainty

Because end of life onset is interval-censored between scheduled visits, assigning the onset to a single time point introduces binning artifacts in survival and hazard estimates. We corrected for this using a fractional binning approach, distributing each individual’s statistical weight across their specific uncertainty interval. For estimators such as Kaplan-Meier and Nelson-Aalen, interval-censored individuals were expanded into multiple fractional copies whose event times were sampled from either a uniform overlap or an empirical conditional distribution restricted to the uncertainty bounds. This strategy preserves the population count while smoothly propagating discrete measurement uncertainty into the continuous estimators. Full mathematical and implementation details are provided in **supplementary note 9**.

For mice data we assumed uniform weight in the uncertainty interval, for humans we assumed uniform weight for the fitting process, and an empirical conditional distribution for plotting. We provide the human plots with uniform weights in **supplementary note 10**.

### Applying observation model to simulation

To directly compare continuous macroscopic damage simulations with discrete empirical longitudinal data, we applied an observation model to the simulated trajectories. For each simulated individual, a longitudinal visit schedule was sampled from empirical data. Simulated Smurf onset was interval-censored to the first scheduled visit following the true threshold crossing, maintaining the interval of uncertainty. Simulated death times were generally right-censored at the study’s end; however, to accurately reflect empirical datasets (where exact death dates are often tracked independently of surveys), a defined constant ratio of individuals retained their exact true death times regardless of the observation window. Left-censored individuals were excluded to prevent bias, and unobserved Smurf onsets preceding a recorded death were imputed to the time of death. Full details of the sampling and constraint logic are provided in **supplementary note 11**.

### Calculation of time from end of life onset to death

In **Fig 2d, Fig 4g-i** and **Fig 5e,f** we present data where the marked dots are the bin centers. The values for each bin are the frequency of individuals in this bin out of the total population. To account for uncertainty in end-of-life onset we used fractional counts as we described here and in **supplementary note 9**.

See attached code for the full implementation.

## Data availability

All raw data can be accessed via the original publications or via the accompanying repository: https://doi.org/10.64898/2026.04.20.719588

## Code availability

All code used to analyze the data as well as creating all the figures is available in https://github.com/NavehRaz/Dynamics_of_the_end_of_life.git.

## Acknowledgements

We thank Kenneth Rockwood and Andrew Rutenberg for their valuable feedback and insightful discussions on this work.

Funding was provided by the European Research Council (grant agreement No. 856487) and by the Sagol Institute for Longevity Research, the Weizmann Institute of Science.

U.A. is the incumbent of the Abisch-Frenkel Professional Chair.

This work was supported in part by the Zuckerman STEM Leadership Program.

This work uses data from the HRS. The Health and Retirement Study (HRS) is supported by the National Institute on Aging (grant number NIA U01AG009740) and is conducted by the University of Michigan.

## Supplementary Information

### 1. Reversibility of end-of-life stage

For Drosophila and mice the original works^19,22^ describe the end-of-life (Smurf) transition as irreversible. From a mathematical perspective, the SR model does not strictly enforce this irreversibility; systemic damage *x* naturally fluctuates and may temporarily drop below the sublethal threshold *X*_*s*_. Indeed, these stochastic drops in systemic damage are responsible for the initial decline in hazard observed post-onset (Main Text **Fig. 6**).

In humans, clinical Frailty Index (FI) scores can exhibit measurement noise and minor natural fluctuations. Therefore, we define the human end-of-life phase as beginning at the *first* observation of *FI* > 0. 55. Once this threshold is crossed, the individual is considered to be in the end-of-life phase until death, agnostic to subsequent reductions in observed FI.

#### Frailty Reversibility Competing-Risk Analysis in humans

Here we analyze the probability that FI improves by different amounts after crossing the threshold we define as end-of-life, using a competing-risk framework. For each observed FI measurement, we treated that measurement as a baseline and followed the individual forward through all subsequent measurements. Reversibility, or “healing,” was defined as the first subsequent FI value that was lower than the baseline by at least a prespecified, whole-deficit threshold.

The healing thresholds were chosen as the nearest whole-deficit equivalents to FI decreases of 0.10, 0.20, 0.30, and 0.35, corresponding to 4 deficits (4/41=0.097), 8 deficits (8/41=0.195), 12 deficits (12/41=0.292), and 14 deficits (14/41=0.341), respectively.

For each baseline observation and threshold, the outcome was classified as one of three mutually exclusive possibilities: healing, death, or censoring. Healing occurred if a later measured FI satisfied:

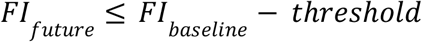

The event time was assigned to the age at the first such measurement. If no such FI decrease was observed before the end of follow-up, individuals were classified as having died or having been censored.

We estimated the cumulative incidence of healing using the Aalen-Johansen estimator, treating death as a competing event and censoring as right-censoring. Thus, the reported reversibility probability is the cumulative incidence of observing a qualifying FI decrease before death, properly accounting for the fact that death precludes later healing.

Estimates were calculated overall and within age-by-baseline-FI strata (**Supplementary Figures S1-S4**). Heatmaps omit baseline-FI strata where the upper edge of the FI bin is less than or equal to the required FI decrease, as reversibility of that magnitude is mathematically impossible.

As demonstrated in the heatmaps, a significant physiological recovery is unlikely. While smaller FI reductions (∼0.1, or 4 deficits) show higher probability (**Fig S1**), the incidence of substantial healing (a reduction of 0.3 in FI, representing recovery from end-stage-frailty to a not frail state) is smaller than 0.1, dropping to near zero for older populations (**Figs S4, S5**).

**Supplementary Figure S1.**
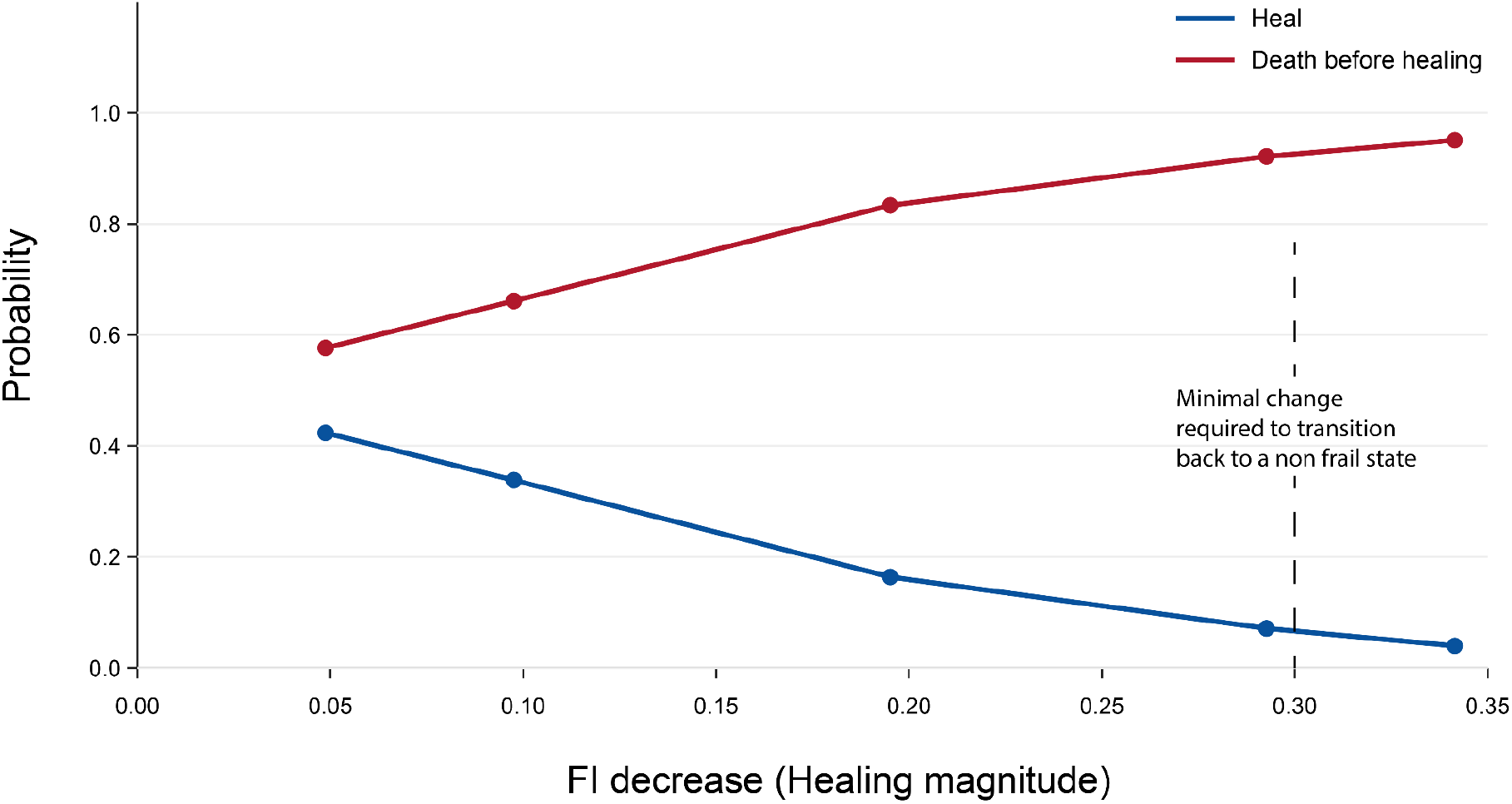
Probability of reversing end stage frailty. The probability to observe a reduction of FI (blue) vs dying (red) for different magnitudes of FI reduction. The probability is calculated on individuals with initial FI of 0.5-0.6. The probability to heal back to a non frail state (FI<0.2) is smaller than 8%.

**Supplementary Figure S2.**
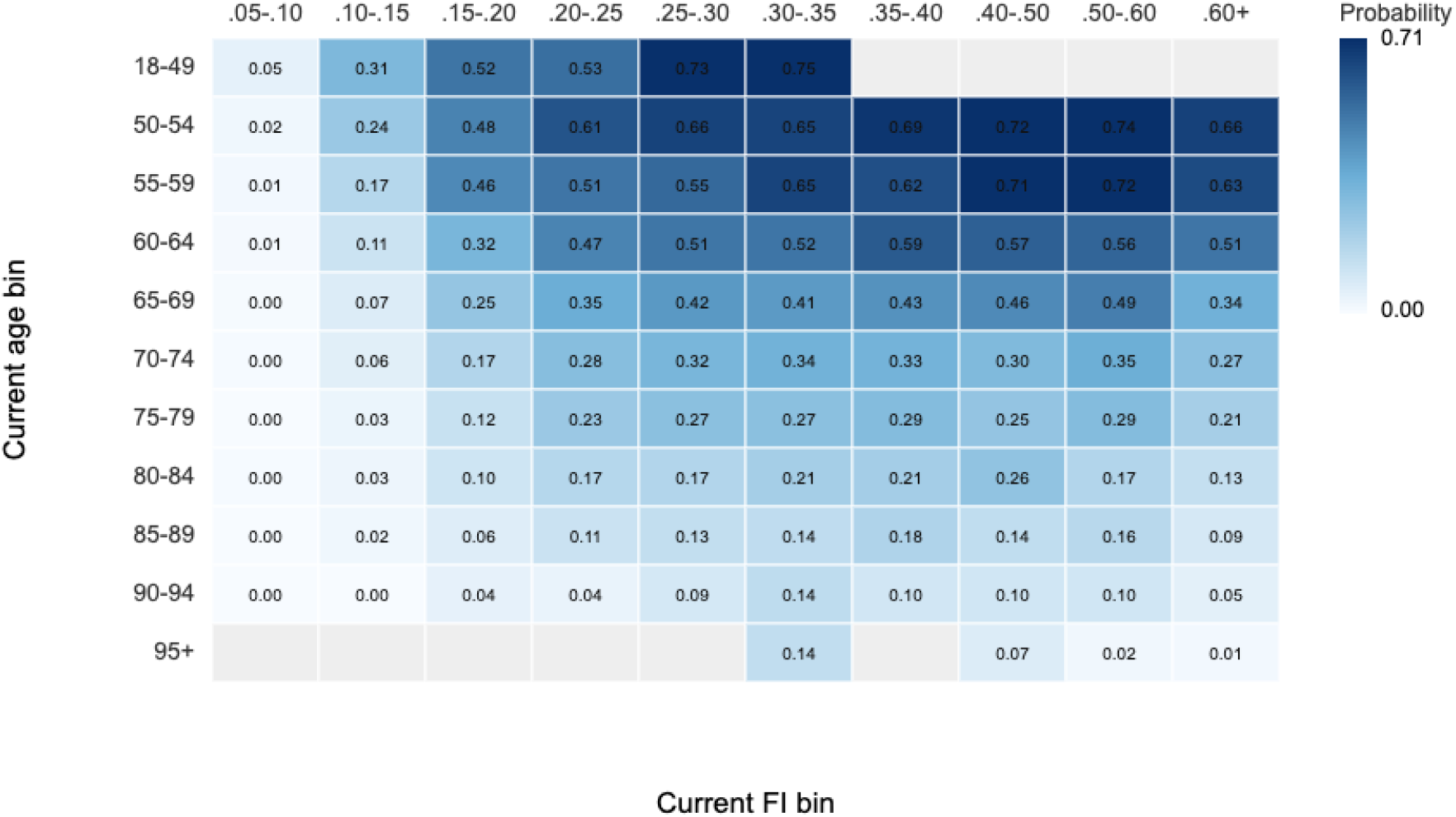
Aelen Johnson estimates for a 4 deficits (∼0.1) reversibility in FI. Grey cells have n<100, values are Aelen Johnson healing CIF, impossibly low FI columns are hidden.

**Supplementary Figure S3.**
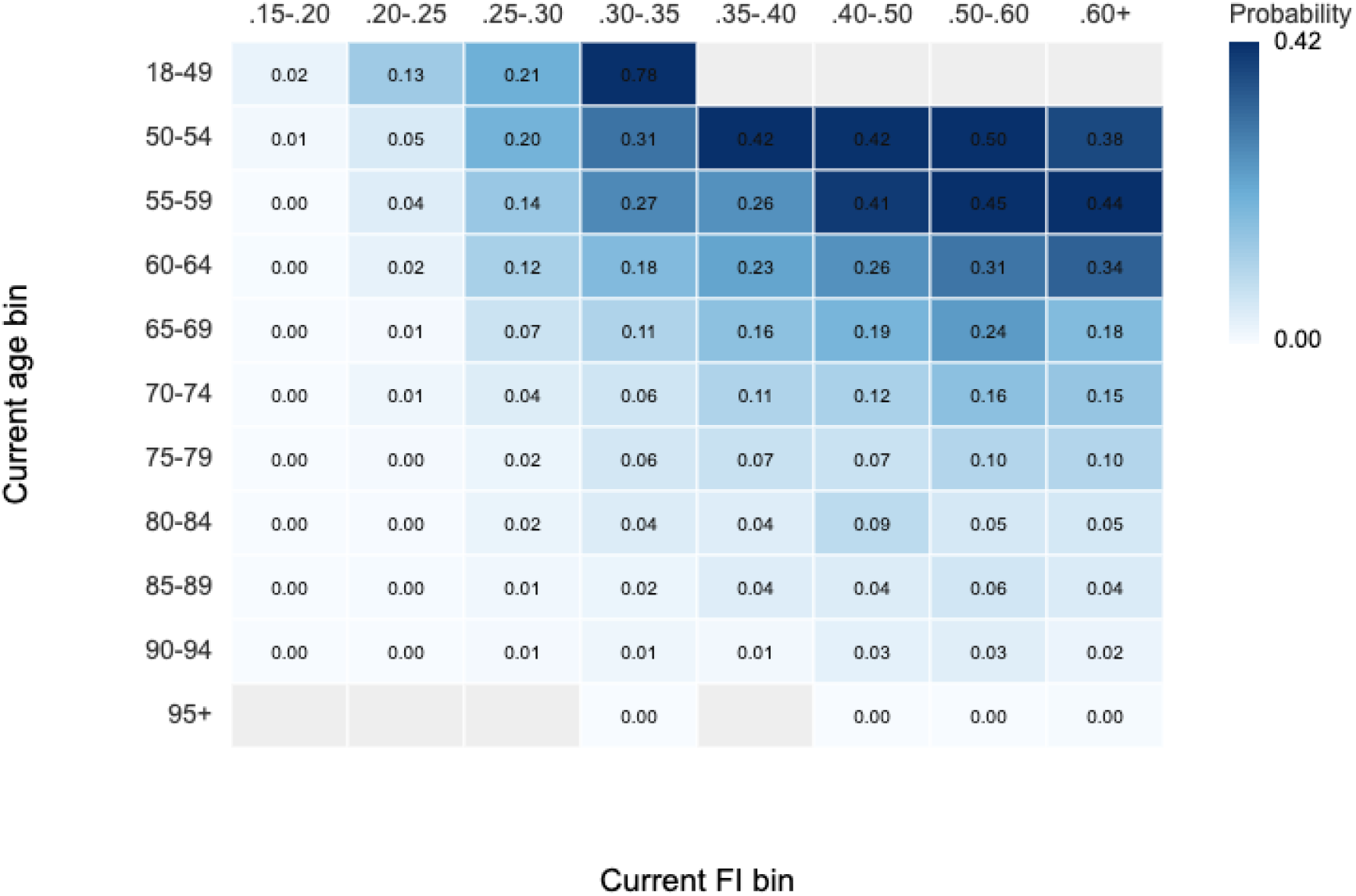
Aelen Johnson estimates for an 8 deficits (∼0.2) reversibility in FI. Grey cells have n<100, values are Aelen Johnson healing CIF, impossibly low FI columns are hidden.

**Supplementary Figure S4.**
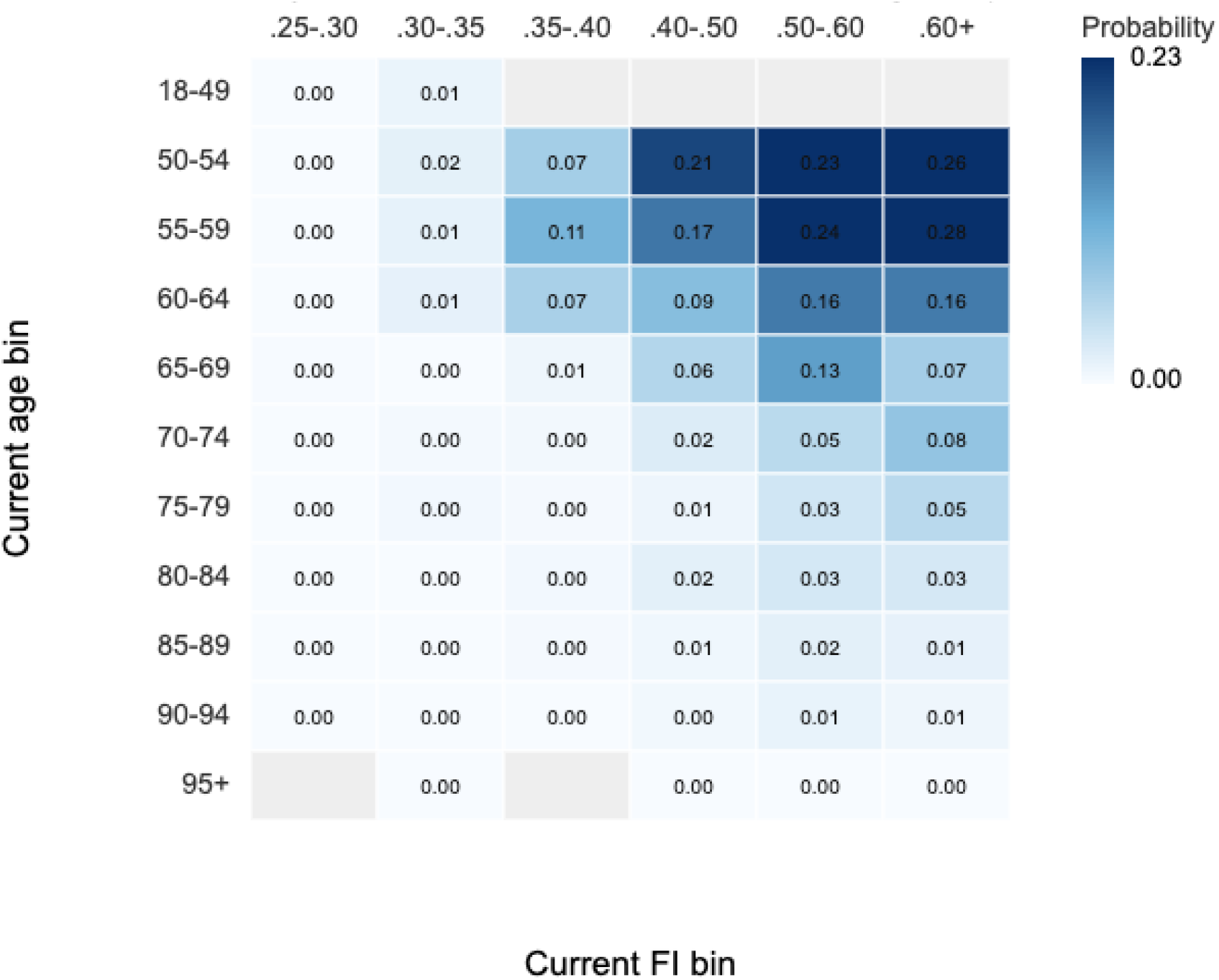
Aelen Johnson estimates for a 12 deficits (∼0.3) reversibility in FI. Grey cells have n<100, values are Aelen Johnson healing CIF, impossibly low FI columns are hidden.

**Supplementary Figure S5.**
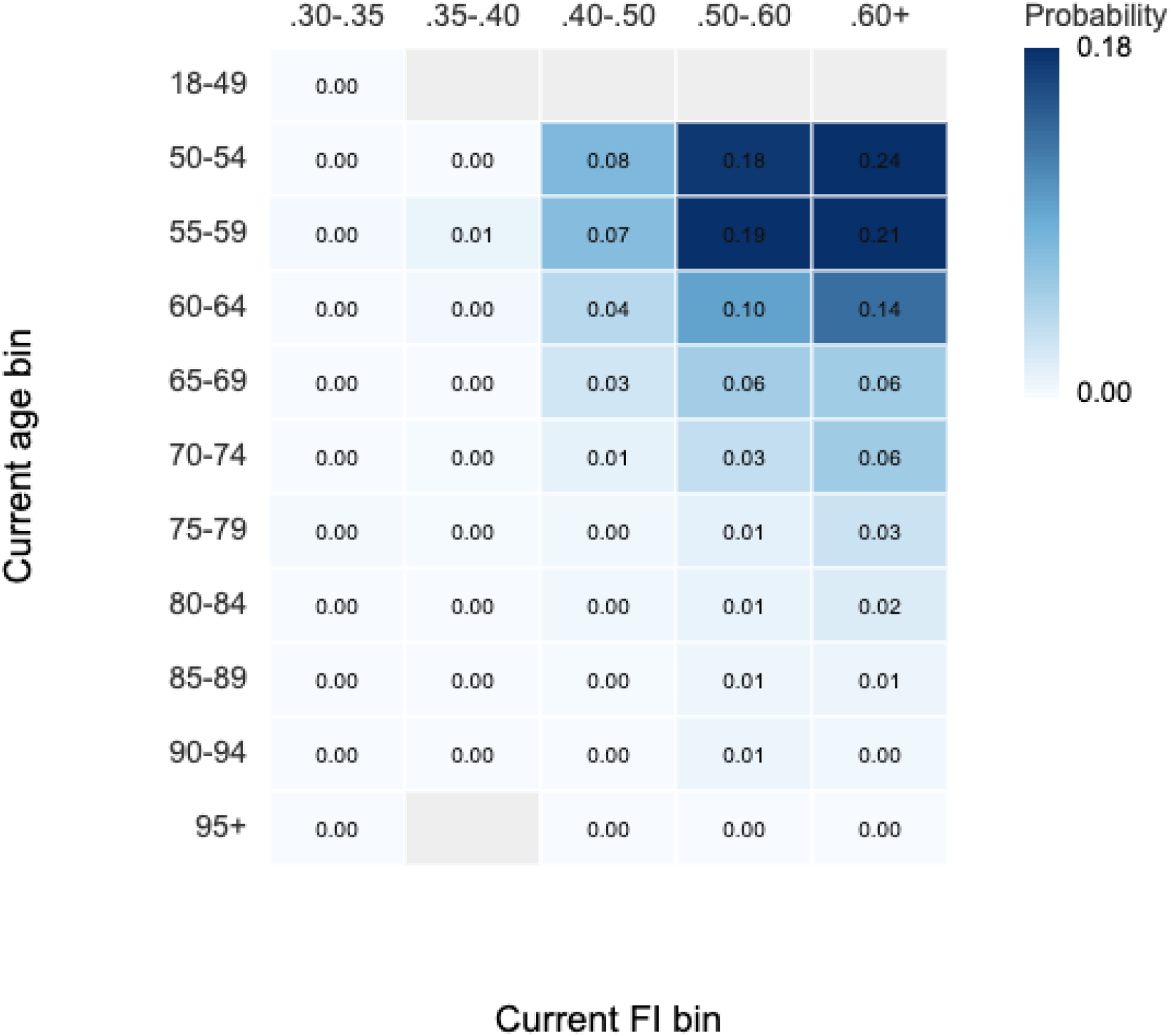
Aelen Johnson estimates for a 14 deficits (∼0.35) reversibility in FI. Grey cells have n<100, values are Aelen Johnson healing CIF, impossibly low FI columns are hidden.

### 2. Median Smurf duration barely change as a function of age of onset

In their original work Tricoire and Rera^19^ observed that the median Smurf duration appears to be independent of the age at which the Smurf transition occurs. In **Supplementary Figure S6**, we show that while the SR model predicts a weak dependence of the median duration on the age of onset, this effect is difficult to resolve in the experimental data as it is obscured by noise.

**Supplementary Figure S6.**
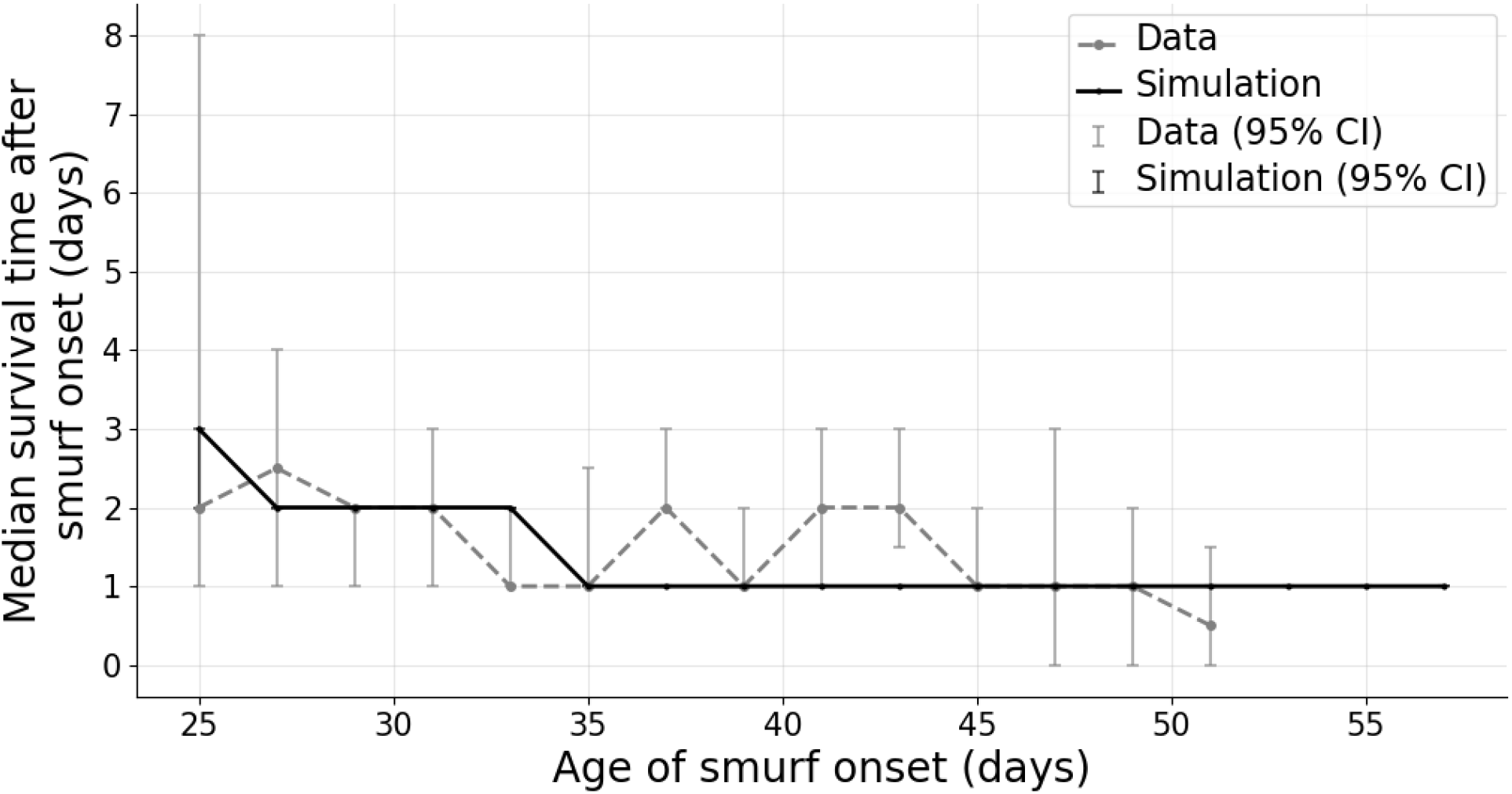
The median Smurf duration is less sensitive to age of Smurf onset. Simulation shows a change from 3 days at 25 days age of onset, 1 day for age of onset >35 days. The data however is more noisy and the 2 days median Smurf duration seems to remain within the CI for most ages.

### 3. Smurf durations for different genotypes

We simulated different *Drosophila melanogaster* genotypes using SR parameters fitted to their survival curves. We find that the model predicts a decline of mean Smurf survival time with onset age in other genotypes as well (**Fig S7c**).

The SR model, calibrated on the survival curves of each fly strain, predicts that the median smurf duration is roughly independent of the strain, about 3-5 days. This is despite a 2-fold difference in the strain median lifespans, between 31 and 55 days (life spans were not strongly correlated with Smurf durations. This prediction agrees with that data (**Fig S7a**). The distribution of Smurf durations is also similar to the data of Tricoire and Rera^19^ (**Fig S7b**).

Due to the data collection methodology for these specific strains, our access was limited to survival data and median Smurf durations. Consequently, we could not directly compare the mean Smurf durations against the empirical data. Median Smurf duration was estimated based on the proportion of Smurfs recorded at each sampling time point.

Median Smurf duration was estimated based on the proportion of Smurfs recorded at each sampling time point. Whereas experimental observations occurred once every 24h, our simulation has exact transition times and so to align our simulation with these experimental observations, we excluded simulated individuals that turned Smurf and died on the same day, as these transient states would not have been captured during the periodic census-taking.

For Genotypes other than drsGFP we had survival data sampled every few days and did not have individual Smurf data. We therefore calibrated for survival using the exact pipeline from^14^ and assumed the same *X*_*s*_/*X*_*c*_ ratio as in the drsGFP flies.

**Supplementary Fig S7.**
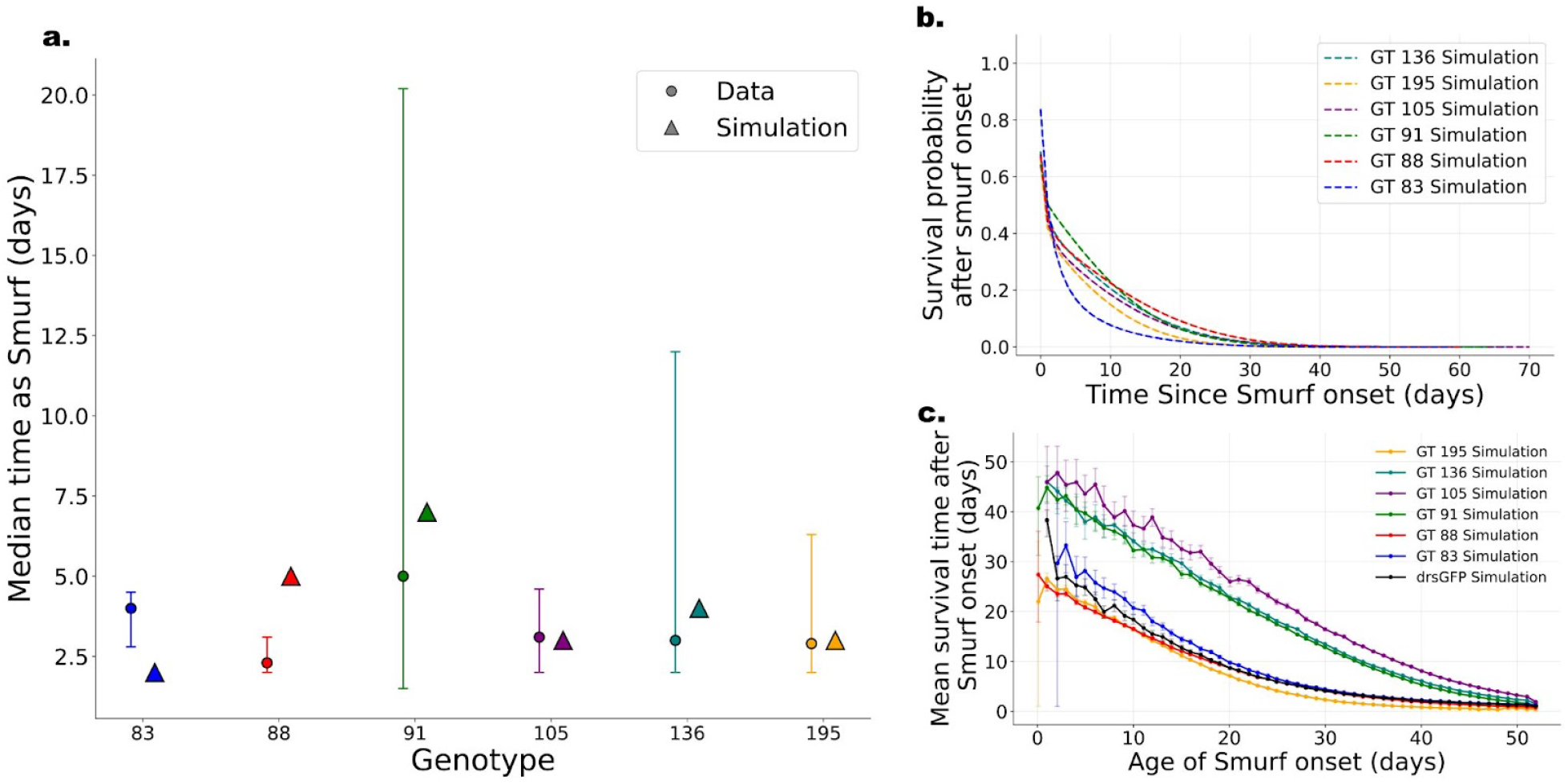
The SR model captures the reduction of Smurf mean duration with onset age, and the similarity of Smurf median duration in different fly strains. **a**. Simulating Smurf conditions for different Drosophila DGRP genotypes generally agrees with previously shown data. Error bars are SEM. **b**. Most genotypes have similar survival probabilities as Smurfs regardless of having very different lifespans, in agreement with previous findings. **c**. Simulation and data show that mean time spent as a Smurf is shorter the later the Smurf onset, in contrast to what was previously described.

### 4. Full fit values and CI

**Supplementary table 1.** Maximum likelihood sets for each dataset. These sets represent the 5 dimensional maximum likelihood and should be used for simulation. These are the parameters used for the plots in the paper.

|  | Humans<br>M | Humans<br>F | Drosophila<br>drsGRP | Mice AKRJ | Mice<br>C57 M | Mice C57 F |
| --- | --- | --- | --- | --- | --- | --- |
| **Production rate**<br>$\eta \left[ \frac{\text{damage}}{\text{day}^2} \right]$ | 7.13e-8 | 1.3e-7 | 0.024 | 8.7e-4 | 0.0031 | 1.1e-4 |
| **Removal rate**<br>$\beta \left[ \frac{\text{damage}}{\text{day}} \right]$ | 0.00295 | 0.006 | 0.84 | 0.155 | 2.86 | 0.092 |
| **Noise magnitude**<br>$\epsilon \left[ \frac{\text{damage}^2}{\text{day}} \right]$ | 1.59e-5 | 2.76e-5 | 0.287 | 0.42 | 3.96 | 0.018 |
| **Death threshold**<br>$X_c [\text{damage}]$ | 1.32 | 1.21 | 4.11 | 19.89 | 33.85 | 5.54 |
| **Smurf threshold as % of death threshold**<br>**Production rate**<br>$X_s [\% X_c]$ | 97.2% | 95.8% | 83.4% | 85.6% | 91.9% | 95.4% |

**Supplementary table 2.**
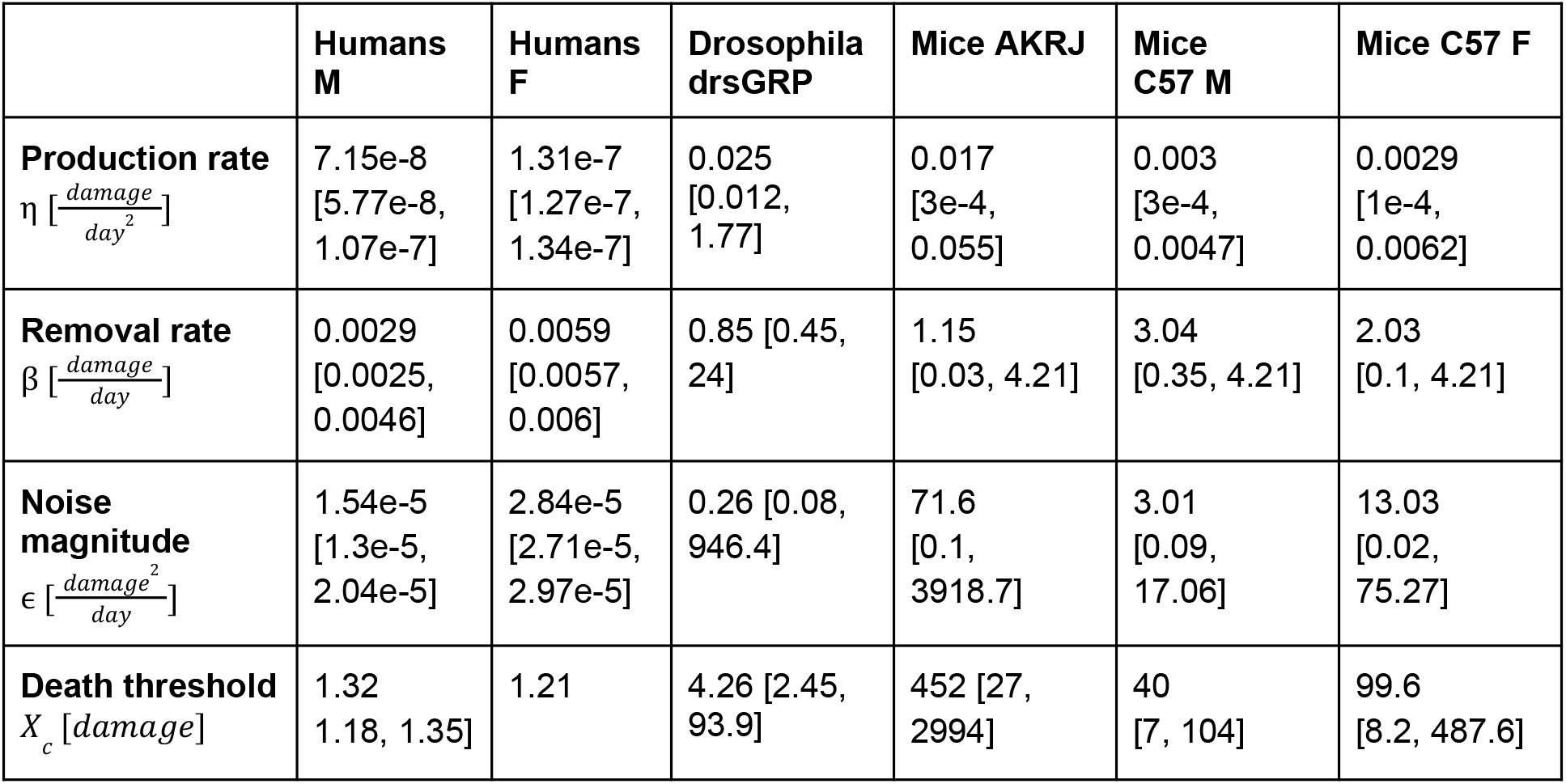
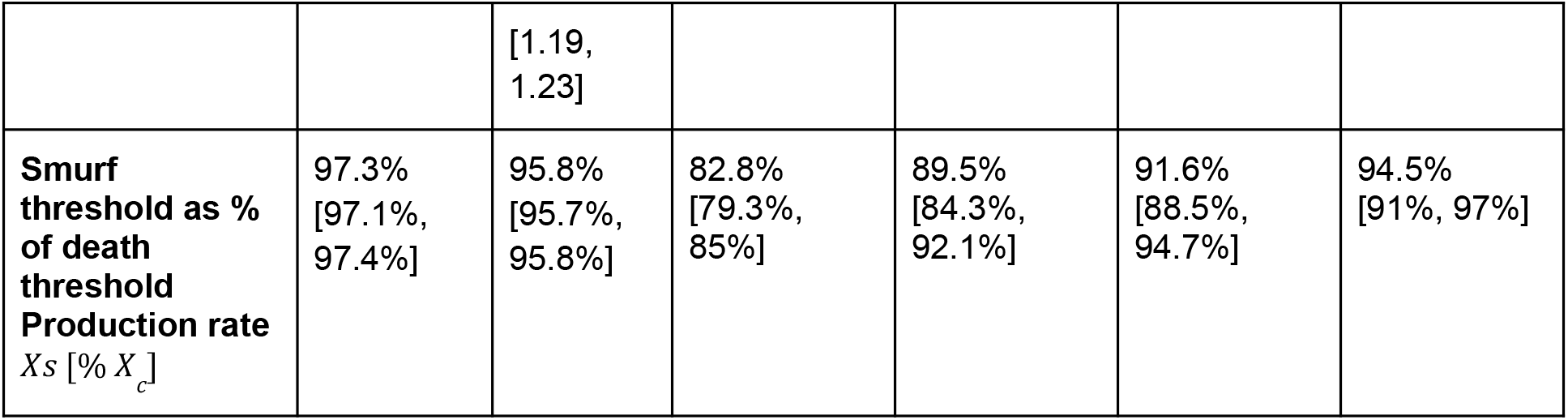
Maximum likelihood of marginalized fit values and 95% CI’s. For each parameter the we integrate over the other parameters and present the maximum likelihood for each individual parameter. This excludes correlations to other params, and these sets should not be used for simulating

### 5. Estimation of End-of-Life Duration as a Function of Onset Age with Censored Data

To quantify how the duration of the end-of-life (Smurf) phase changes depending on the age at which it begins, individuals are grouped into age bins based on their observed end of life onset time. To account for the interval censoring of the onset time itself, the fractional binning method (see **SIXX**) is utilized to distribute an individual’s statistical weight across overlapping age bins.

Within each age bin, estimating the average duration strictly from individuals with observed deaths would introduce a survivorship bias, as right-censored individuals (who survived past the observation window) would be improperly excluded. To resolve this, a Kaplan-Meier (KM) survival estimator is fit to the post-Smurf durations (the time elapsed from Smurf onset to either death or censoring) for all individuals occupying the bin.

Summary statistics for the duration are then derived directly from the censorship-aware KM curve:

- **Median Duration:** Extracted directly as the median survival time from the bin-specific KM fit.
- **Mean Duration:** Estimated using the Restricted Mean Survival Time (RMST). To ensure robust and comparable estimates across all age bins, the RMST is computed by integrating the area under the bin’s KM survival curve up to a fixed, global time horizon (the maximum observed post-Smurf duration across the entire dataset).

To quantify statistical uncertainty, 95% confidence intervals are generated using a bootstrapping approach (e.g., 1000 iterations). In each iteration, the data within the bin (or the fractional weights) are resampled with replacement, the KM curve is refit, and the median or RMST is re-calculated, yielding confidence bounds that incorporate both sampling variance and measurement uncertainty.

In **Supplementary Figure S8** we show the mean end-of-life duration as a function of onset age for humans with and without the observation model vs the data. In all cases there is a clear “shortening twilight” where the mean end-of-life duration is shorter the later the onset. Except for young ages, the data seems to be between the raw simulation to the simulation in the observation model, suggesting that perhaps a better observation model will fully capture the data.

The discrepancy at younger ages might be due to

**Supplementary Figure S8.**
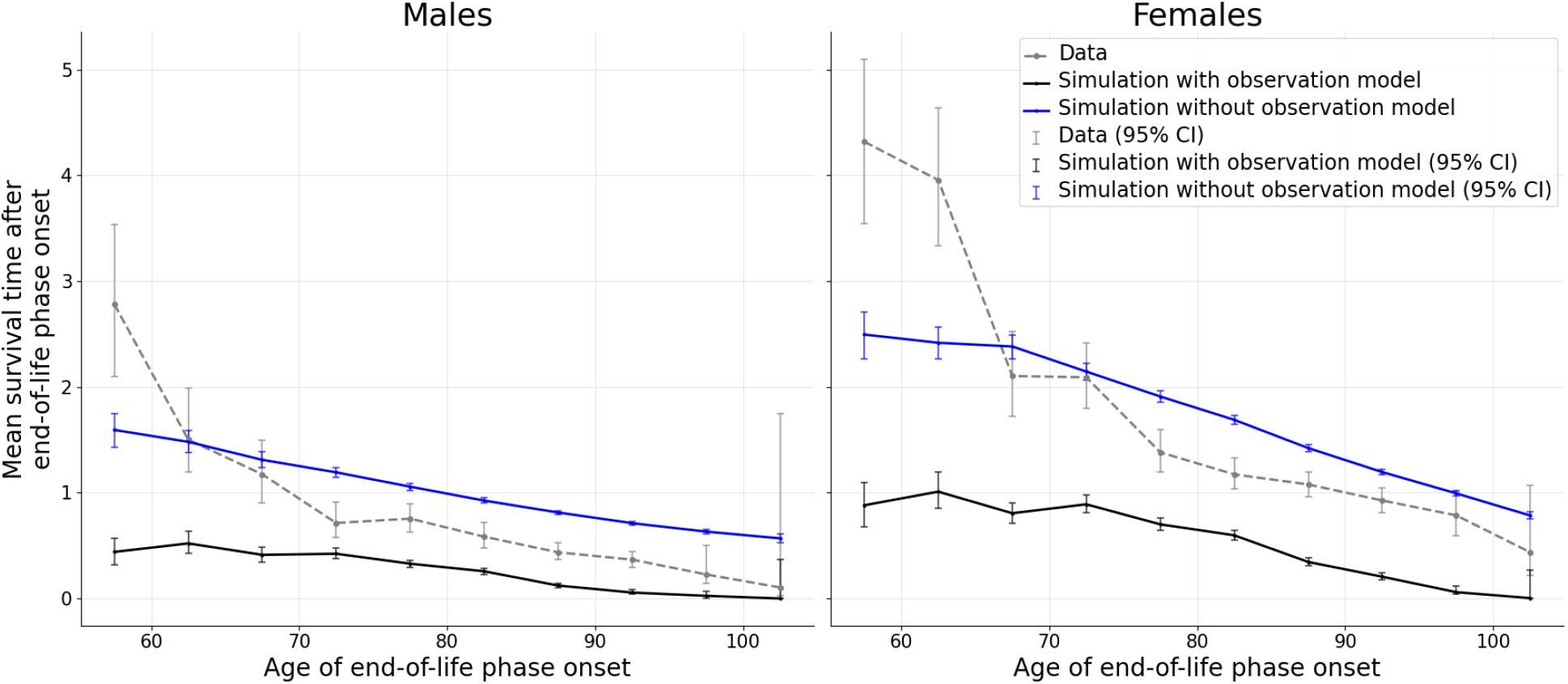
Human mean end of life duration as a function of age of onset. Blue lines are simulations without observation model, black lines are simulation with observation model, data is in gray dashed line.

### 6. Likelihood Estimation for Joint Death and Smurf-Duration Distributions

To evaluate the agreement between experimental observations and stochastic simulations, we employ a **Dirichlet-multinomial (DM) likelihood** framework. This approach accounts for the joint distribution of death times (*T*_*d*_) and end-of-life (Smurf) durations (*τ* = *T*_*d*_ − *T*_*s*_), while incorporating experimental measurement uncertainty and right-censoring.

#### 1. Data Binning and Interval Uncertainty

The 2D state space is discretized into a histogram with a bin size of Δt. We define the valid domain such that *τ* ≤ *T*_*d*_, corresponding to the physical constraint that an end-of-life transition cannot occur before birth (*T*_*s*_≥ 0).

A key feature of this implementation is the handling of **interval uncertainty** in experimental data. Experimental observations are often recorded at discrete intervals (dt). Rather than treating an observation as a single point, we distribute its probability mass across the 2D bins (*T*_*d*_, *τ*) overlapped by its uncertainty window. This results in “fractional counts” *y*_*i*_ for each bin *i*, ensuring the likelihood is not biased by the arbitrary placement of bin edges.

#### 2. Dirichlet-Multinomial Framework

The simulation generates exact event times, which are binned to provide counts *c*_*i*_. To account for finite-sample stochasticity and avoid zero-probability issues, the simulation defines the parameters (pseudo-counts) of a Dirichlet distribution: *α*_*i*_ = *c*_*i*_ + λ where λ = 0.5 represents the **Jeffreys prior**. The total log-likelihood for uncensored observations, In *L*_*u*_, is:

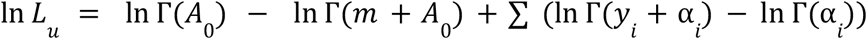

Where:

- *m* is the total count of valid experimental events.
- *A*_0_ = ∑ *α*_*i*_ is the sum of the prior strengths across all valid bins (sum of pseudo-counts).
- Γ is the Gamma function.

#### 3. Handling of Censored Data

For right-censored observations (where an individual dies before end-of-life transition, or the experiment concludes prior to death), we incorporate their contribution In *L*_*c*_ by integrating the model’s probability mass over the region of the state space compatible with the censoring constraints:

- **Death-censored:** If a end-of-life transition is observed at *T*_*s*_ but the individual is alive at *T*_*censor*_, the likelihood includes the sum of probabilities for all bins where *T*_*d*_ > *T*_*censor*_ and *τ* > *T*_*censor*_ − *T*_*s*_.
- **Both-censored:** If no transition is observed by *T*_*censor*_, we sum the probabilities for all bins where *T*_*s*_ > *T*_*censor*_.

The total log-likelihood is given by: In *L*_*total*_ = In *L*_*u*_ + In *L*_*c*_

### 7. Data preparation

Upon the introduction of bromophenol blue dye on day 12, the experimental hazard rate exhibits a transient elevation that deviates from the long-term, monotonic, aging trend until approximately day 24 (**Supplementary Figure S9**). This “acclimation” period likely reflects extrinsic mortality associated with the transfer process or physiological adjustment to the dye-labeled medium. We excluded this period from the fitting process to ensure the model accurately captures the dominant intrinsic aging process rather than transient experimental effects.

**Supplementary Figure S9.**
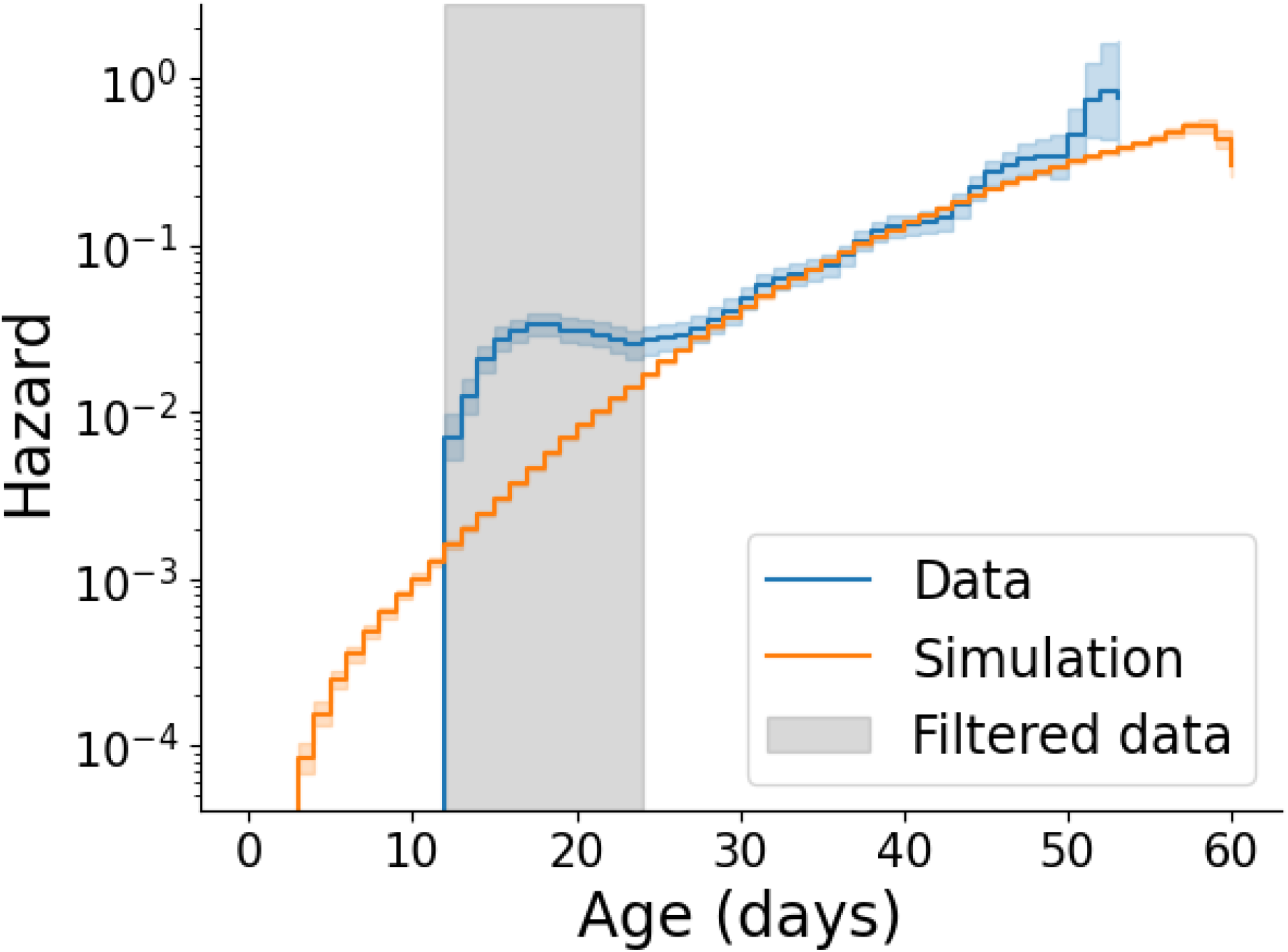
The experimental drosophila hazard rate (blue) exceeds the modeled trajectory (orange) between days 12 and 24. This period (shaded gray) was excluded from the likelihood calculation to ensure the model accurately captures the dominant intrinsic aging process rather than transient experimental effects (such as extrinsic death).

### 5. Health variables used for frailty index (HRS)

**Table S1.** Health variables used for frailty index (HRS).

| Number | Variable | Description | Encoding |
| --- | --- | --- | --- |
| 1 | armsa | Difficulty reaching/extending arms up | 0: no, 1: yes/any |
| 2 | arthre | Ever diagnosed with arthritis | 0: no, 1: yes |
| 3 | batha | Difficulty bathing or showering | 0: no, 1: yes/any |
| 4 | bathh | Gets help bathing/showering | 0: no, 1: yes |
| 5 | beda | Difficulty getting in/out of bed | 0: no, 1: yes/any |
| 6 | bede | Uses equipment to get in/out of bed | 0: no, 1: yes |
| 7 | bedh | Gets help getting in/out of bed | 0: no, 1: yes |
| 8 | cancre | Ever diagnosed with cancer | 0: no, 1: yes |
| 9 | chaira | Difficulty getting up from chair | 0: no, 1: yes/any |
| 10 | clim1a | Difficulty climbing one stair flight | 0: no, 1: yes/any |
| 11 | climsa | Difficulty climbing several stair flight | 0: no, 1: yes/any |
| 12 | diabe | Ever diagnosed with diabetes | 0: no, 1: yes |
| 13 | dimea | Difficulty picking up a dime | 0: no, 1: yes/any |
| 14 | dressa | Difficulty dressing | 0: no, 1: yes/any |
| 15 | dressh | Gets help dressing | 0: no, 1: yes |
| 16 | eata | Difficulty eating | 0: no, 1: yes/any |
| 17 | eath | Gets help eating | 0: no, 1: yes |
| 18 | hearte | Ever diagnosed with heart problems | 0: no, 1: yes |
| 19 | hibpe | Ever diagnosed with high blood pressure | 0: no, 1: yes |
| 20 | homcar | Received home health care within previous 2 years | 0: no, 1: yes |
| 21 | hosp | Had a hospital stay within previous 2 years | 0: no, 1: yes |
| 22 | lifta | Difficulty lifting/carrying 10lbs | 0: no, 1: yes/any |
| 23 | lunge | Ever diagnosed with lung disease | 0: no, 1: yes |
| 24 | moneya | Difficulty managing money | 0: no, 1: yes/any |
| 25 | nhmliv | Living in nursing home at time of interview | 0: no, 1: yes |
| 26 | nrshom | Had a nursing home stay within previous 2 years | 0: no, 1: yes |
| 27 | outpt | Had outpatient surgery within previous 2 years | 0: no, 1: yes |
| 28 | phonea | Difficulty using the telephone | 0: no, 1: yes/any |
| 29 | pusha | Difficulty pushing/pulling a large object | 0: no, 1: yes/any |
| 30 | shlt | Self-reported health | 0: excellent-good, 1: fair-poor |
| 31 | shopa | Difficulty shopping for groceries | 0: no, 1: yes/any |
| 32 | spcfac | Visited a specialized health facility within previous 2 years | 0: no, 1: yes |
| 33 | stoopa | Difficulty stooping/kneeling/crouching | 0: no, 1: yes/any |
| 34 | stroke | Ever diagnosed with a stroke | 0: no, 1: yes |
| 35 | toilta | Difficulty using the toilet | 0: no, 1: yes/any |
| 36 | toilth | Gets help using the toilet | 0: no, 1: yes |
| 37 | walk1a | Difficulty walking one block | 0: no, 1: yes/any |
| 38 | walkra | Difficulty walking across rooms | 0: no, 1: yes/any |
| 39 | walkre | Needs equipment to walk across rooms | 0: no, 1: yes |
| 40 | walkrh | Gets help walking across rooms | 0: no, 1: yes |
| 41 | walksa | Difficulty walking several blocks | 0: no, 1: yes/any |

### 9. Fractional Binning for Interval-Censored Data

Because end of life onset is evaluated at discrete intervals, the exact time of onset is inherently unknown and falls within an interval *I*_*i*_ = [*t*_*obs,i*_ − *Δt*_*i*_, *t*_*obs,i*_]. Assigning the onset simply to the time of observation or the interval midpoint can introduce binning artifacts and biases in survival and hazard estimates. To correct for this, a fractional binning logic distributes each individual’s statistical weight across their specific uncertainty interval.

The unit mass of each interval-censored individual *i* is distributed across overlapping age bins *B*_*k*_ = [*b*_*k*_, *b*_*k*+1_) using one of two strategies:

1. **Uniform Overlap Strategy:** The individual’s mass is spread uniformly across the interval. The fractional weight *w*_*i,k*_ added to age bin *B*_*k*_ is exactly proportional to the temporal overlap:

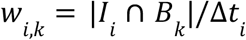
2. **Empirical Age-Bin Spreader:** To account for non-uniform probabilities of onset within an interval, mass can be distributed using an empirical conditional distribution generated from the simulation. Simulated individuals are grouped into buckets using a known anchor variable *A*_*i*_ (e.g., the exact death time for individuals with observed deaths). The fractional weight assigned to each bin is derived from the empirical distribution of true simulated end of life times (*T*_*S*_) within the matching anchor bucket, restricted exclusively to the [*t*_*obs,i*_ − *Δt*_*i*_, *t*_*obs,i*_] bounds:

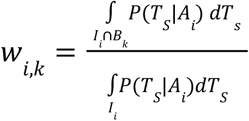

If an anchor bucket contains too few simulated samples to form a reliable distribution, the method automatically falls back to the uniform overlap strategy.

#### Application to Statistical Estimators

To seamlessly incorporate fractional binning into standard estimators such as Kaplan Meier used for survival estimation and Nelson Aalen used for hazard estimations, each interval-censored individual is expanded into *n* fractional copies with exact time of event bootstrapped from the probability (uniform or empirical spreader) and typically *n = 10* or *5*. Each copy is assigned a fractional weight of *1/n*. The event times for these copies are sampled directly from the empirical conditional distribution across the uncertainty interval. This weighted expansion preserves the overall population count while propagating the discrete measurement uncertainty into the final calculations.

### 10. Fits without observation model and with uniform overlap fractional binning

As detailed in methods and **supplementary notes 9 and 11**, to generate the plots for humans in the main text we used an observation model and an empirical age-bin spreader that relies on the simulated distribution to weigh the data in the measurement uncertainty interval.

In **Supplementary Figure 10** we show plots of the simulation without the observation model vs the data with uniform fractional binning.

**Supplementary Figure S10.**
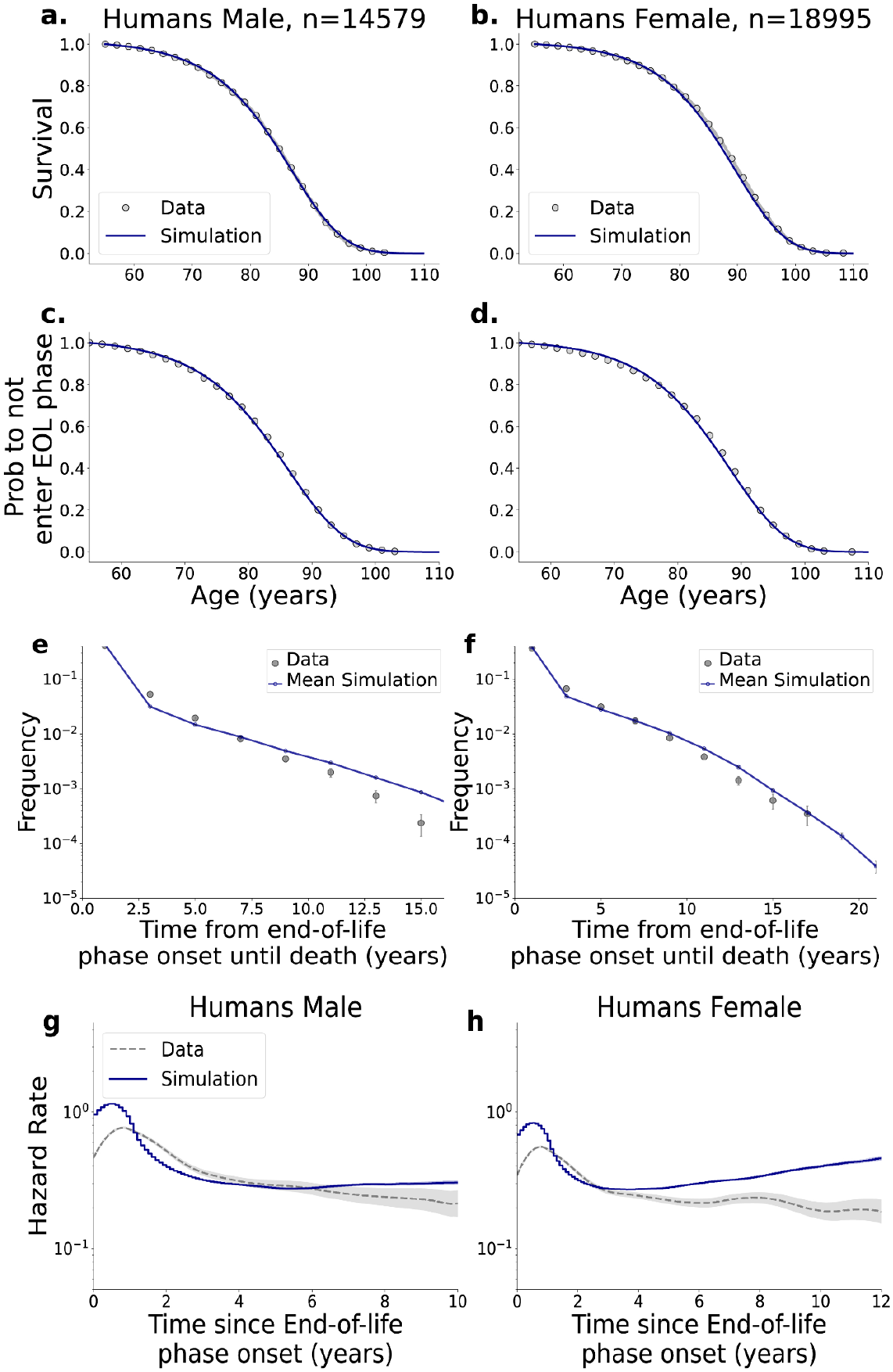
Human plots without observation model and with uniform binning.

### 11. Application of observation model to simulated data

To directly compare continuous macroscopic damage simulations to the discrete longitudinal measurements of empirical datasets with limited study duration, specifically the human bi-yearly HRS surveys, an observation model is applied to the simulated trajectories. This process transforms exact, continuous simulated times into interval-censored observations that mirror empirical sampling constraints. For each simulated individual, a longitudinal visit schedule *V* = {*v*_1_, *v*_2_, ···, *v_n_*} and an exact-death tracking flag are sampled with replacement from the empirical observation times. The continuous simulated times for end of life onset (*T*_*S*_) and death (*T*_*D*_) are mapped to this schedule using the following logic:

1. **Exact Death Tracking vs. Right-Censoring:** Empirical datasets often track exact death dates for a subset of individuals independently of the survey window (e.g., via national registries). If the sampled empirical flag dictates keeping the exact death time, the observed death time is the true simulated death time (*TD,obs = TD*). Otherwise, death is right-censored at the final scheduled visit: *T*_*D,obs*_ = min (*T*_*D*_, *v*_*n*_). This ensures the simulation maintains the exact constant ratio of uncensored-to-censored deaths present in the empirical data.
2. **Interval-Censored EOL Onset:** If a simulated individual crosses the end of life threshold, the onset is only recorded at the first scheduled visit following the true simulated crossing: *T*_*s, obs*_ = *v*_*j*_ where *v*_*j*_ = min (*v* ∈ *V*|*v* > *T*_*s*_) The interval of uncertainty (*Δt*) is strictly tracked as the duration between this positive visit and the immediately preceding visit:

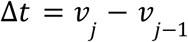

#### Imputation at Death

If a simulated individual dies before the next scheduled visit can record the end of life onset, the EOL onset is imputed to coincide exactly with the time of death:

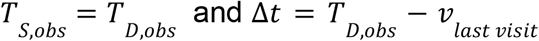

### 12. Human hazard during end-of-life phase as a function of age of onset

In **Figure S11**, we show the post-end-of-life phase onset hazard for humans, filtered by 20-year age groups. In **Figure S12**, we present a similar analysis, instead allowing for individuals older than a specific minimum age threshold (filtering out individuals younger than the specified age, as previously described).

**Supplementary Fig S11.**
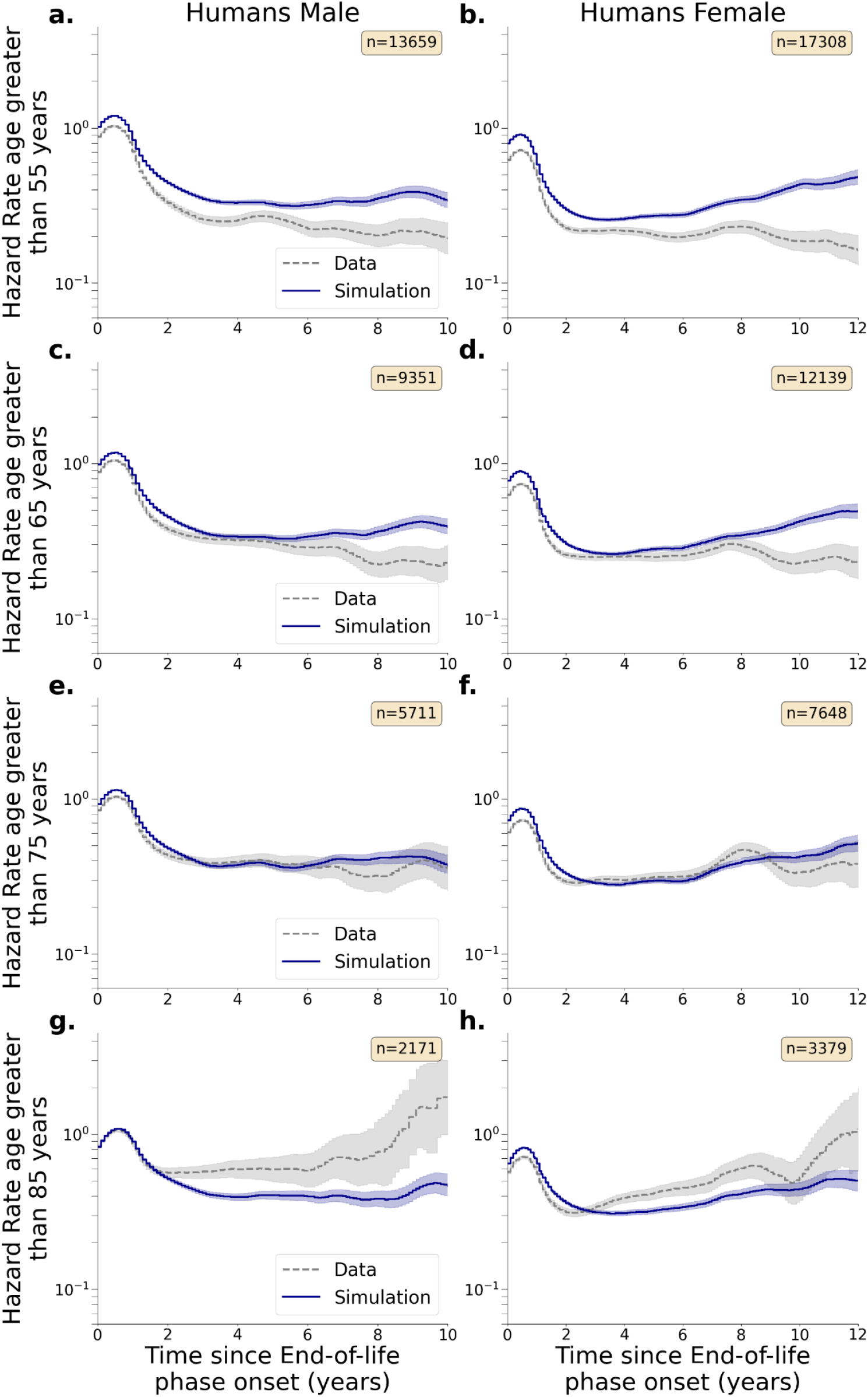
Human Hazard post end-of-life filtered by age groups. **a**,**b -** hazard for individuals with age of onset between 55-75 years. **c**,**d**. Hazard for individuals with age of onset between 65-75 years. **e**,**f**. Hazard for individuals with age of onset between 75-95 years. **g**,**h**. Hazard for individuals with age of onset between 85-105 years

**Supplementary Fig S12.**
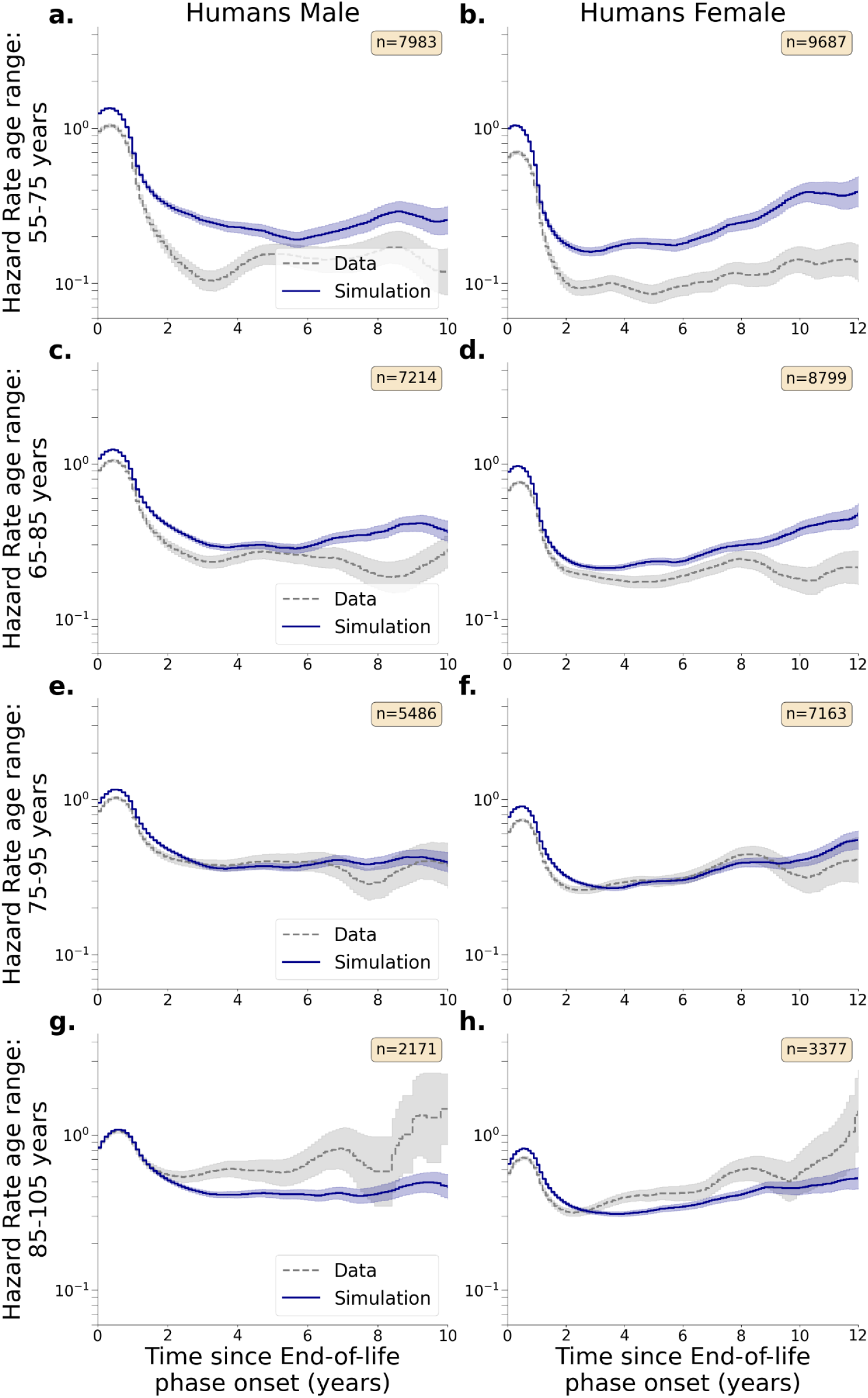
Human Hazard post end-of-life filtered by minimal onset age. **a**,**b -** hazard for individuals with age of onset greater than 55 years. **c**,**d**. Hazard for individuals with age of onset greater than 65 years. **e**,**f**. Hazard for individuals with age of onset greater than 75 years. **g**,**h**. Hazard for individuals with age of onset greater than 85 years.

### 13. Effect of changing the FI threshold in humans

We tested the effects of changing the end-of-life onset FI threshold in humans.

#### Impact on Data Distributions

We chose to exclude left censored individuals to avoid introducing biases to the fitting process. Since the left censoring probability decreased the higher the threshold, there was a minor effect on the survival curve (**Fig S13 a,b)**.

Changing the FI threshold also significantly changed the distribution of the end-of-life onset and its duration (**Fig S13 c-h)**.

#### Impact on Goodness of Fit

Modifying the FI threshold resulted in a lower likelihood for the model fits as detailed in **Table S2**. The fit quality is also visually poorer as can be seen in **Fig S14, Fig S15**.

**Table S2.** goodness of fit as a function of FI threshold: the table shows likelihood normalized to the sample size for the relevant threshold (sample sized differed because we avoided left censoring)

| FI threshold | Log Likelihood per individual (females) | Log Likelihood per individual (males) |
| --- | --- | --- |
| 0.55 | -1.95 | -2.02 |
| 0.2 | -2.47 | -2.63 |
| 0.1 | -2.51 | -2.78 |

**Supplementary Fig S13.**
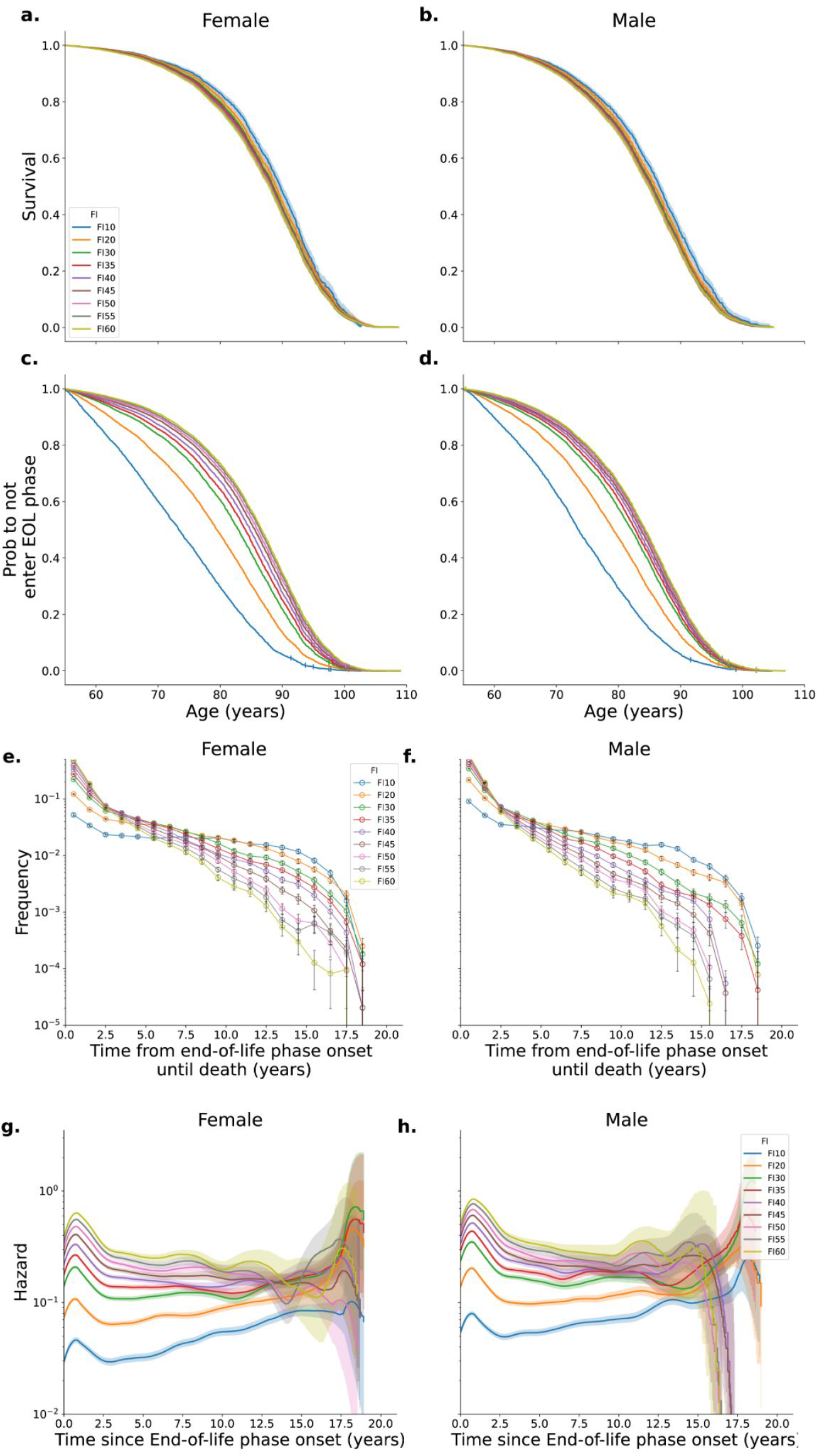
Effects of changing the FI threshold on the end-of-life and survival. **a**,**b**. Survival values for different FI thresholds. **c**,**d**. Probability to not enter the end-of-life phase as a function of age for different FI thresholds. **e**,**f**. Duration of the end-of-life phase until death for different FI thresholds. **g**,**h**. Hazard in the end-of-life phase for different FI thresholds

**Supplementary Fig S14.**
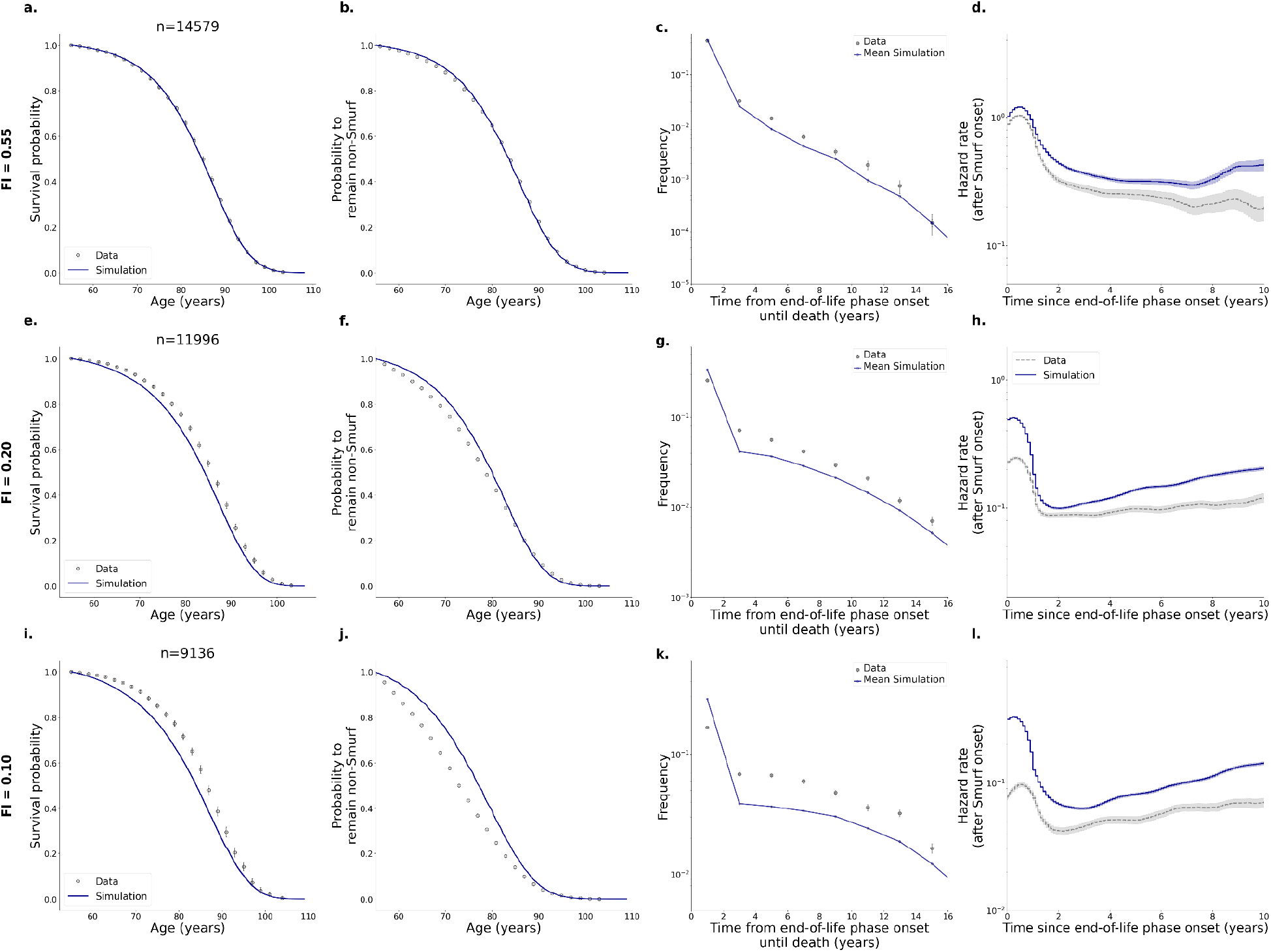
Low FI threshold results in poorer fits for men. **a-d**. FI =0.55. **e-h**. FI=0.2, **i-l**. FI =0.1.

**Supplementary Fig S15.**
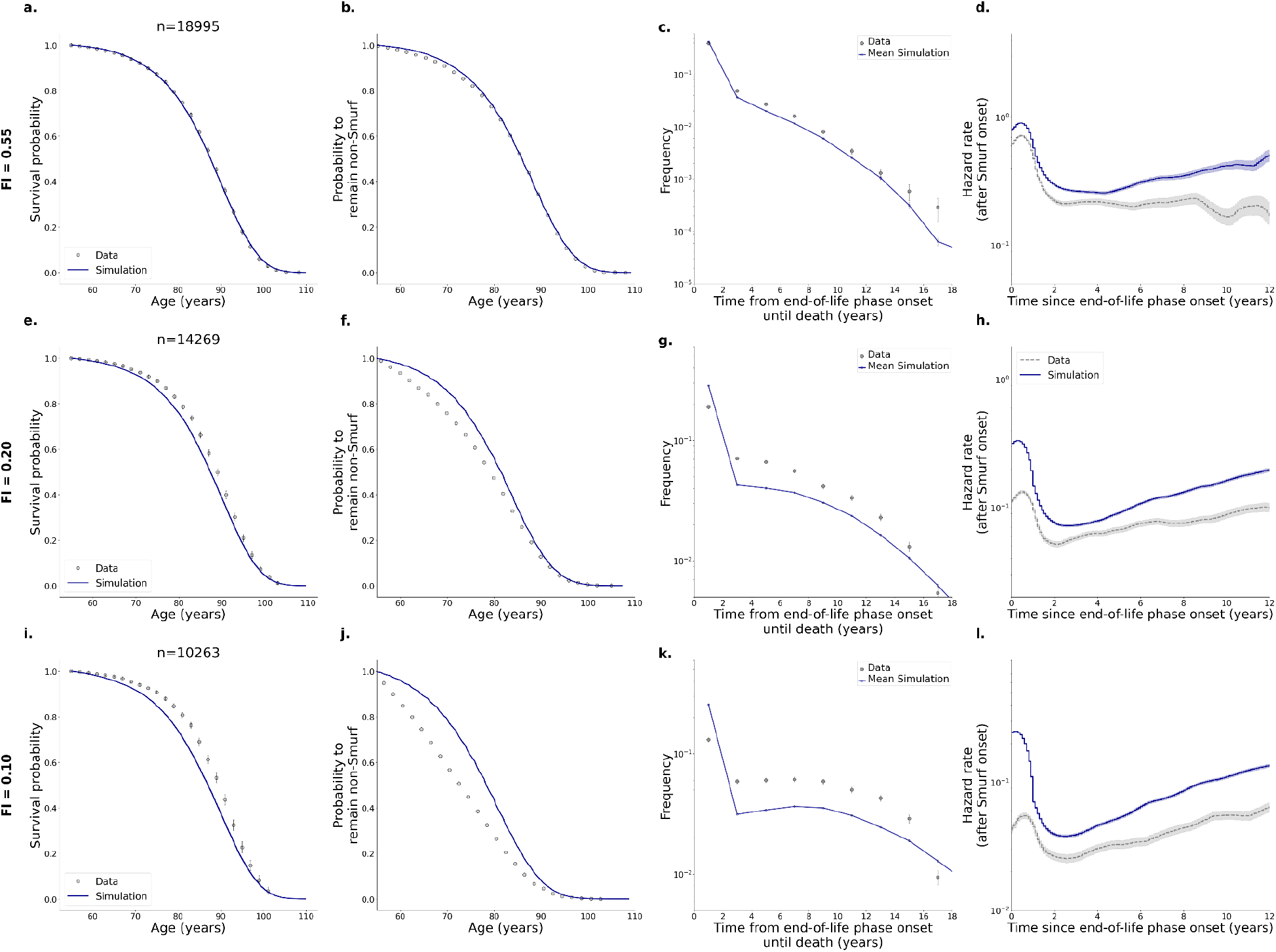
Low FI threshold results in poorer fits for women. **a-d.** FI =0.55. **e-h**. FI=0.2, **i-l**. FI =0.1.

### 14. Randomizing end-of life onset time smoothes the end-of-life hazard

To confirm that the initial reduction in hazard after end-of-life onset is a biological feature and not an artifact of our plotting methodology, we randomized the end-of-life onset times for the human data while keeping the exact death times intact. This randomization resulted in the complete disappearance of the initial drop in hazard, validating our original observation.

**Supplementary Fig S16.**
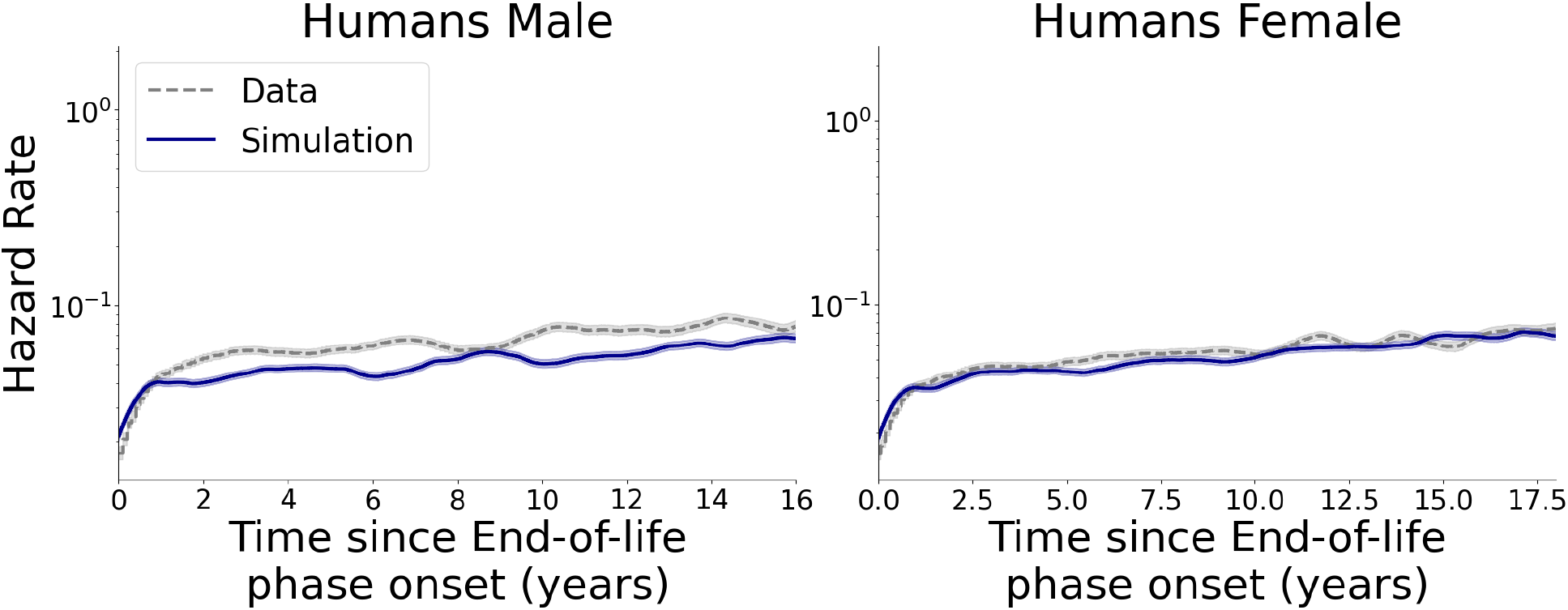
initial drop in hazard is non-existent when Randomizing end-of-life onset times.

### 15. Dataset sizes, filtering and imputations

**Table S3.** Number of individuals in each dataset and how many were removed at each part of the data processing. Imputed individuals are individuals where death was observed before the end-of-life transition and the transition time was assumed to occur between the last observation and the time of death (see supplementary note 9).

| Organism | Total N | Time range applied | N in time range | N -left censored (removed ) | N - used | N - right censored in time range | N imputed In time range |
| --- | --- | --- | --- | --- | --- | --- | --- |
| **Drosophila** | 1159 | 25-55 days | 817 | 149 | 668 | - | - |
| **Mice AKR/J** | 45 | - | 45 | - | 45 | - | 8 |
| **Mice C57BL6/J males** | 44 | - | 44 | - | 44 | - | 16 |
| **Mice C57BL6/J females** | 48 | - | 48 | - | 48 | - | 24 |
| **Human males** | 14952 | 55-110 years | 14036 | 361 | 13675 | 8532 | 4228 |
| **Human females** | 19720 | 55-110 years | 18027 | 686 | 17340 | 11539 | 4162 |

